# The Essential Role of Latrophilin-1 Adhesion GPCR Nanoclusters in Inhibitory Synapses

**DOI:** 10.1101/2023.10.08.561368

**Authors:** Daniel Matúš, Jaybree M. Lopez, Richard C. Sando, Thomas C. Südhof

## Abstract

Latrophilin-1 (Lphn1, a.k.a. CIRL1 and CL1; gene symbol *Adgrl1*) is an Adhesion GPCR that has been implicated in excitatory synaptic transmission as a candidate receptor for α-latrotoxin. Here we analyzed conditional knockin/knockout mice for Lphn1 that contain an extracellular myc-epitope tag. Surprisingly, we found that Lphn1 is localized in cultured neurons to synaptic nanoclusters that are present in both excitatory and inhibitory synapses. Conditional deletion of Lphn1 in cultured neurons failed to elicit a detectable impairment in excitatory synapses but produced a decrease in inhibitory synapse numbers and synaptic transmission that was most pronounced for synapses close to the neuronal soma. No changes in axonal or dendritic outgrowth or branching were observed. Our data indicate that Lphn1 is among the few postsynaptic adhesion molecules that are present in both excitatory and inhibitory synapses and that Lphn1 by itself is not essential for excitatory synaptic transmission but contributes to inhibitory synaptic connections.

## INTRODUCTION

Synapses are the fundamental unit of neuronal networks that mediate sensation, cognition and behavior. An enormous body of work described many of the fundamental processes that underlie synapse function, such as postsynaptic neurotransmitter reception, presynaptic vesicle release, or various forms of synaptic plasticity (reviewed in Kandel 2001, Südhof 2013, Basu and Siegelbaum 2015, Bailey et al. 2015, Chen and Gouaux 2019, Scott and Aricescu 2019). However, the mechanisms and molecules that enable synapses to be formed in the first place are poorly understood. Recent insights into synapse formation came from the study of synaptic adhesion molecules, such as neurexins, neuroligins, or adhesion GPCRs (aGPCRs) (reviewed in Südhof 2017, Cao and Tabuchi 2017, Südhof 2018, Sanes and Zipursky 2020, Gomez et al. 2021, Fuccillo and Pak 2021, Boxer and Aoto 2022, Cortés et al. 2023). Adhesion GPCRs are characterized by a combination of a large extracellular N-terminal region containing a variety of adhesion domains, a so-called GAIN-domain that mediates autoproteolysis and is invariably found in Adhesion GPCRs, a canonical 7 transmembrane domain typical for all GPCRs, and a rather long cytoplasmic region (reviewed in Hamann et al. 2015, Vizurraga et al. 2020, Rosa et al. 2021, Einspahr and Tilley 2022, Liebscher et al. 2022, Seufert et al. 2023). This domain combination places aGPCRs at the intersection of cell-cell or cell-matrix adhesion and signal transduction, although the precise functions of most aGPCRs are unclear. Multiple aGPCRs, including brain angiogenesis inhibitors (BAI’s), latrophilins, and CELSR’s, are thought to contribute to synapse formation (Najarro et al. 2012, Sigoillot et al. 2015, Wang et al. 2021, Wang et al. 2020, Bolliger et al. 2011, Shiu et al. 2022, Tu et al. 2018, Zhu et al. 2015, Stephenson et al. 2013, Aimi et al. 2023, Martinelli et al. 2016, Kakegawa et al. 2015, Freitas et al. 2023, Li et al. 2022, Zhou et al. 2021, Thakar et al. 2017, Anderson et al. 2017, Sando et al. 2019), although limited information is available about their mechanisms of action and scope of functions.

Three latrophilins are expressed in mice (protein names Lphn1-3; gene symbols *Adgrl1-3*) (Sugita et al. 1998, Ichtchenko et al. 1999), with primary transcripts that are subject to extensive alternative splicing (Sugita et al. 1998, Boucard et al. 2014, Ovando-Zambrano et. al. 2019). Latrophilins are postsynaptic receptors (Anderson et al. 2017, Sando et al. 2019) that contain an N-terminal lectin-like and olfactomedin-like domains that bind to presynaptic teneurin (Silva et al. 2011, Boucard et al. 2014) and presynaptic Flrt adhesion molecules (O’Sullivan et al. 2012), respectively. Intracellularly, latrophilins contain long cytoplasmic sequences that are expressed in multiple alternatively spliced variants, of which the most prevalent variant interacts with intracellular SHANK proteins (Kreienkamp et al. 2000, Tobaben et al. 2000), which are a component of postsynaptic density-protein scaffolds (Naisbitt et al. 1999). Super-resolution microscopy of the latrophilin ligand teneurin-3 revealed that it is not ubiquitously distributed within a synapse but assembled into nanoclusters (Zhang et al. 2022). Interestingly, latrophilin-mediated synapse formation requires G-protein coupling (Sando and Südhof 2021), suggesting that synaptic adhesion complexes constitute sites of active signaling. In the hippocampus, Lphn2 and Lphn3 mediate excitatory synapse formation onto different subcellular regions of CA1 pyramidal neurons independent of one another (Sando et al. 2019), whereas in the cerebellum Lphn2/3 act redundantly in parallel-fiber synapse formation (Zhang et al. 2020). Such “molecular codes” are also employed by other synaptic adhesion molecules, such as neurexins or neuroligins (summarized in Südhof 2017, Cao and Tabuchi 2017, Südhof 2018; Sanes and Zipursky 2020, Gomez et al. 2021, Fuccillo and Pak 2021, Boxer and Aoto 2022, Cortés et al. 2023). It is essential to investigate how each synaptic adhesion molecule contributes to synapse formation in order to generate a comprehensive view of the intricate complexity of neuronal connections in the brain.

Previous studies on latrophilins have been focused on Lphn2 and Lphn3, while Lphn1 has not been studied as intensely. Progress in studying Lphn1 was hindered by the lack of specific antibodies and behavioral abnormalities of Lphn1 constitutive KO mice that impeded breeding (Tobaben et al. 2002, Vitobello et al. 2022). A recent study revealed that human loss-of-function mutations of the Lphn1 ortholog (*ADGRL1*) cause major neurodevelopmental impairments and that a mouse mutant with a constitutive loss of Lphn1 exhibits major synaptic impairments (Vitobello et al. 2022). To further investigate the role of Lphn1 in brain, we here generated conditional knockout (cKO) mice that also allow localization of the endogenous protein by virtue of a knocked-in myc epitope tag. We show using STED microscopy that Lphn1 forms nanoclusters in both excitatory and inhibitory synapses of hippocampal neurons. However, in contrast to the other two isoforms, Lphn1 specifically mediates inhibitory synapse formation onto the cell soma. Neuronal morphology was unchanged after Lphn1 deletion. These findings expand our knowledge of how latrophilins define different forms of synaptic connections, be it in excitatory versus inhibitory synapses or in the spatial restriction of synapses to subcellular regions of target neurons.

## RESULTS

### Generation and validation of Lphn1 conditional knockin/knockout (cKO) mice

Earlier studies of Lphn2 and Lphn3 were facilitated by the availability of conditional knockin/knockout mice in which endogenous Lphn2 or Lphn3 was tagged with an epitope for immunocyto-/immunohistochemical localization but could be deleted by expression of Cre recombinase (Anderson et al. 2017, Sando et al. 2019). Although constitutive Lphn1 KO mice were described (Tobaben et al. 2002, Vitobello et al. 2022), breeding Lphn1 KO mice is challenging because of a major behavioral phenotype (Tobaben et al. 2002, Vitobello et al. 2022). Moreover, no highly specific Lphn1 antibodies are available. To overcome these challenges, we generated new Lphn1 conditional knockin/knockout (cKO) mice in which endogenous Lphn1 is tagged at the N-terminus with a myc epitope but can be deleted with Cre recombinase (Figure 1A).

**Figure 1:**
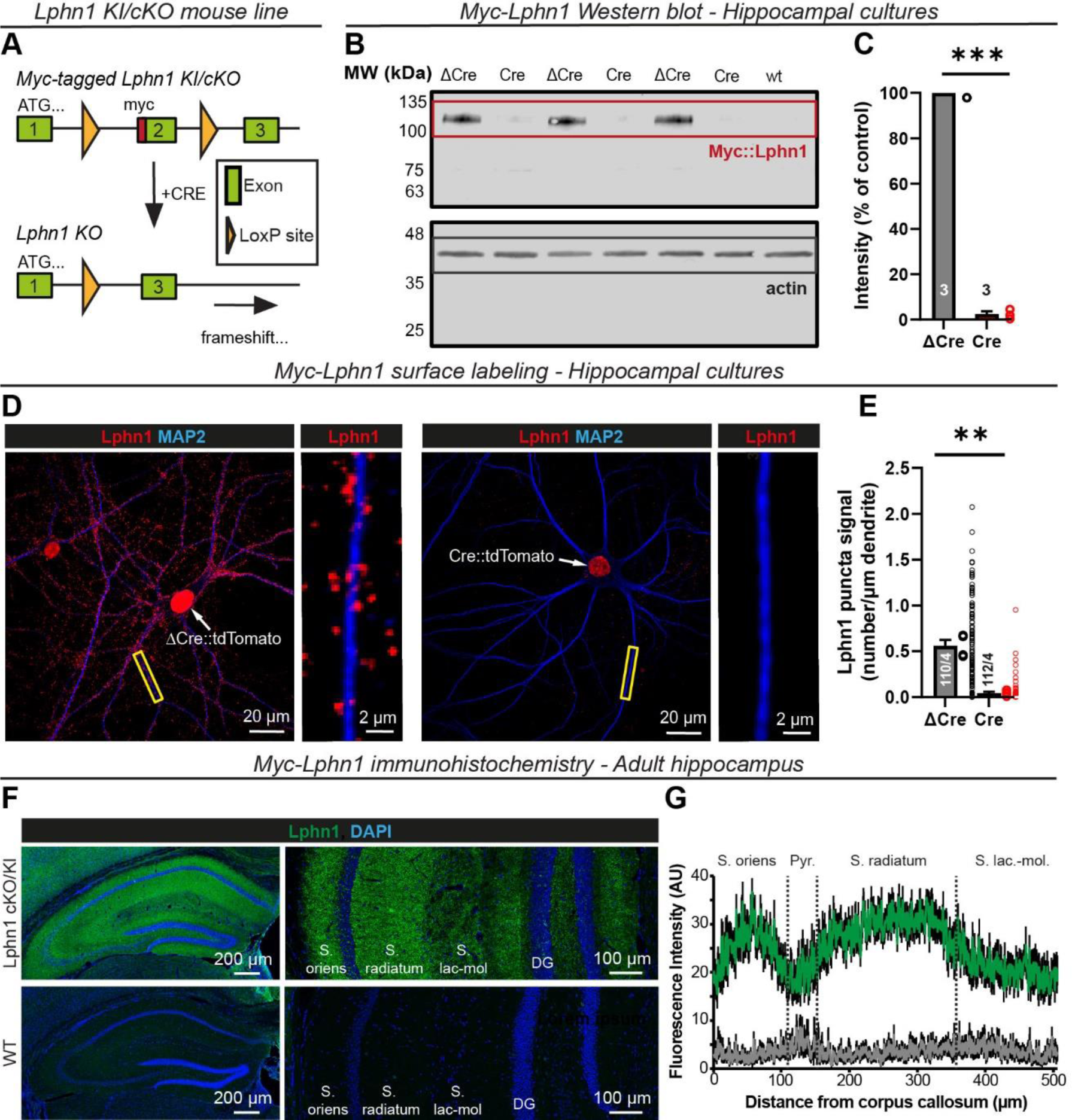
Generation of Latrophilin-1 (Lphn1) conditional knockout (cKO) mice in which endogenous Lphn1 carries an N-terminal myc-tag. (A) Design of Lphn1 cKO mice. A 2xmyc tag sequence was introduced into the second exon of the Lphn1 gene (Adgrl1) and the exon was flanked by loxP sites using homologous recombination in ES cells. (B) Immunoblotting validation of Lphn1 cKO mice. Three independent primary hippocampal cultures from Lphn1 cKO mice were infected with lentiviruses encoding active (Cre) or enzymatically inactive recombinases (ΔCre) that contained a nuclear localization signal and were fused to tdTomato. In addition, hippocampal cells from a single wild-type culture (wt) were analyzed. Blots were stained for the knocked-in myc-tag (top) or actin as a loading control (bottom). Note that owing to GAIN-domain mediated autocleavage, the myc-tag labels the N-terminal Lphn1 fragment of ∼115 kDa instead of uncleaved Lphn1 at ∼210 kDa. Full blot images are shown in Supplementary Figure 1. (C) Quantification of the myc-Lphn1 signal in (B). Signals after subtraction of the wt background were normalized to actin and the ΔCre condition (means ± SEMs; n = 3 independent cultures; statistics were performed using a paired t-test, with ***, p < 0.001). (D) Surface labeling of myc-tagged Lphn1 in primary hippocampal cultures from Lphn1 cKO mice, infected with lentiviruses as described in (B). After myc-staining and permeabilization, neurons were additionally stained for MAP2 to visualize dendrites. Myc-Lphn1 staining shows punctate structures concentrated at dendrites that are largely absent in the Cre condition. The nuclear red fluorescence is derived from the expressed Cre- or ΔCre-tdTomato fusion proteins. (E) Quantification of myc-Lphn1 puncta per µm of dendrite as shown in (D) (means ± SEMs; large dots = mean values for independent cultures; small dots = individual dendritic values; n = 110 dendrites/3 independent cultures; statistics were performed using a paired t-test, with **, p < 0.01). (F) Immunohistochemistry of cryosections from adult Lphn1 cKO and wild-type control mice stained for the myc epitope, showing that myc-Lphn1 is expressed in all layers of the murine hippocampus. (G) Quantification of the myc-Lphn1 fluorescence intensity of stained brain slices from (F) as a function of the CA1-region hippocampal layer (n = 5 (control) or 7 (Lphn1 cKO); green = Lphn1 cKO mice carrying a myc epitope tag; gray = wild-type control).

Lphn1 cKO mice were viable and fertile with no apparent behavioral abnormality. To validate the Lphn1 myc-tagging and conditional Lphn1 deletion, we cultured primary hippocampal cells from newborn mice and infected them with lentiviruses expressing inactive mutant ΔCre recombinase (as a control) or active Cre recombinase (test) under control of the neuronal synapsin-1 promoter. Both active Cre and mutant inactive ΔCre recombinase were expressed as tdTomato fusion proteins that contain a nuclear localization signal which translocates the Cre and ΔCre proteins into the nucleus. Upon expression of Cre but not ΔCre, the floxed exon in Lphn1 cKO neurons is recombined, leading to a frameshift in the mRNA that abolishes Lphn1 protein synthesis. To test the efficiency of Cre recombination, we analyzed total protein from infected cultures by quantitative immunoblotting (Fig 1B, Suppl. Figure 1A). Whereas ΔCre virus-infected cultures exhibited a ∼115 kDa band corresponding to the Lphn1 N-terminal fragment that results from physiological autocleavage by Lphn1’s GAIN domain (Araç et al. 2012), cultures infected with Cre viruses showed a >97% reduction of Lphn1 protein expression (Figure 1B, C). Thus, Cre expression abolished Lphn1 expression in the cultured cells.

We next performed surface labeling of the myc-Lphn1 protein in the same hippocampal cultures (Figure 1D). We observed punctate myc signals along dendrites in the ΔCre condition that were largely absent from neurons in the Cre condition (Figure 1D, E), consistent with the Lphn1 protein deletion. Furthermore, when we stained cryosections of the brains from adult mice for myc-Lphn1 protein, we found that myc-Lphn1 was uniformly distributed among the various strata of the hippocampus (Figure 1F, G), indicating that the Lphn1 distribution differs from that of its paralogs Lphn2 and Lphn3 that are highly enriched in the *S. lacunosum-moleculare* (Lphn2) or the *S. oriens* and *S. radiatum* (Lphn3) (Anderson et al. 2017, Sando et al. 2019).

### Lphn1 forms nanoclusters in both excitatory and inhibitory synapses

Lphn2 and Lphn3 are essential for excitatory but not inhibitory synapse formation and are localized to excitatory synapses, although their possible presence in inhibitory synapses has not been investigated (Anderson et al. 2017, Sando et al. 2019). We therefore examined whether Lphn1 localizes to excitatory and/or inhibitory synapses. As an initial approach, we surface-labelled myc-tagged Lphn1 in primary hippocampal cultures from the Lphn1 knockin/cKO mice, followed by detergent permeabilization of the cells and staining for excitatory and inhibitory synaptic markers (Figure 2, 3). We observed prominent co-labeling of myc-Lphn1 with synaptic markers but found that, surprisingly, Lphn1 co-localized with both excitatory (Figure 2A) and inhibitory synaptic markers (Figure 3A) in the cultured neurons.

**Figure 2:**
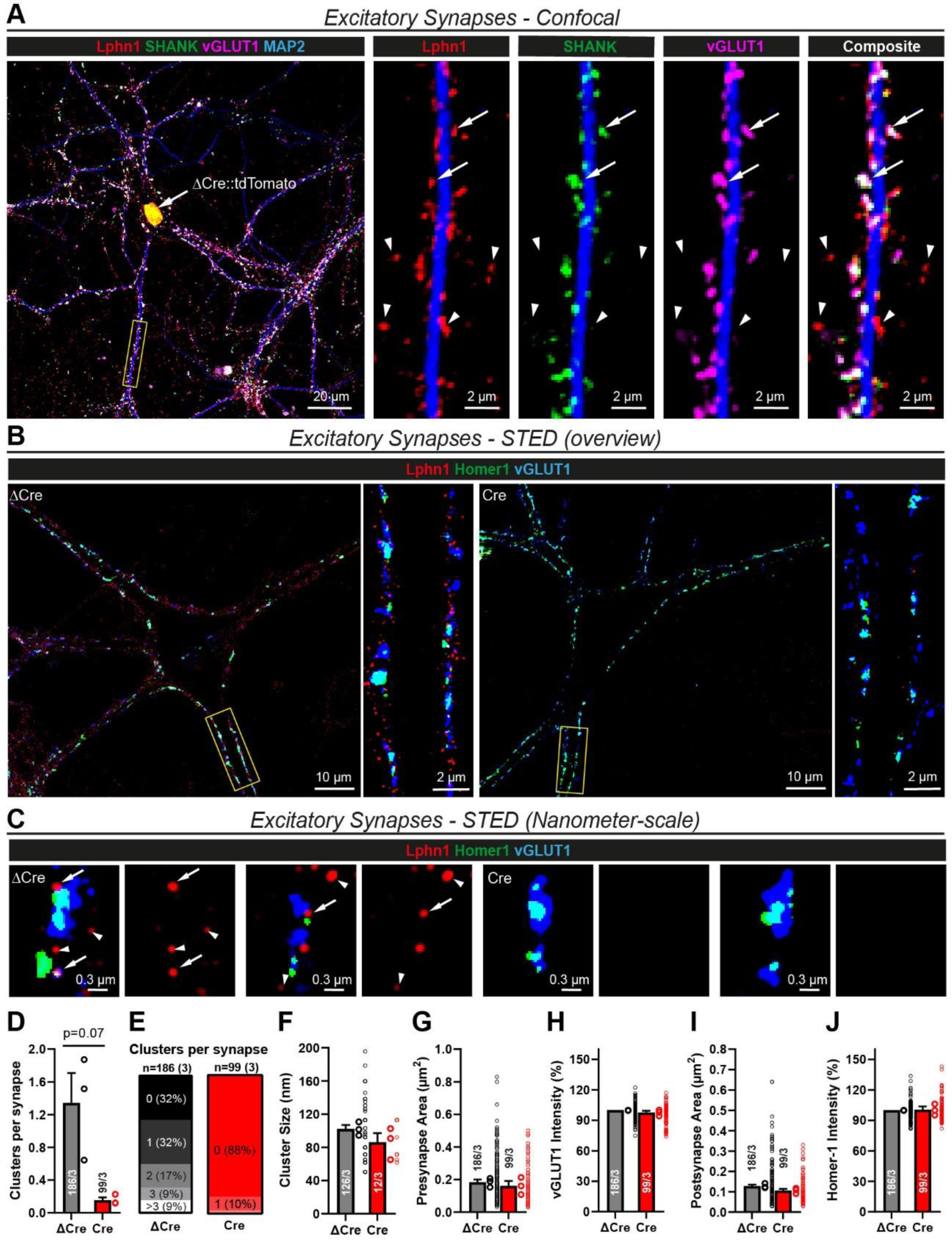
Lphn1 forms nanoclusters in excitatory synapses. (A) Co-localization of myc-Lphn1 with excitatory synapse markers using standard diffraction-limited confocal microscopy. After surface labeling for myc-Lphn1, Lphn1 cKO hippocampal cultures were permeabilized and stained for presynaptic (vGLUT1) and postsynaptic (SHANK) markers. Lphn1 is expressed in excitatory synapses (arrows), but also shows Lphn1 localization to puncta that are not stained for excitatory synapse markers (arrow heads). (B, C) STED microscopy of Lphn1 cKO hippocampal cultures reveals nanoclusters of Lphn1 in excitatory synapses. Cultures were infected with lentiviruses encoding active (Cre) or mutant inactive Cre-recombinase (ΔCre), surface labelled for myc-Lphn1, permeabilized, and stained for presynaptic (vGLUT1) and postsynaptic (Homer1) markers of excitatory synapses. Subsets of Lphn1 nanoclusters co-localize (arrows) or do not co-localize with excitatory synapse markers (arrowheads). Almost no nanoclusters can be found on neurons expressing Cre recombinase. Images are shown at different scales: an overview with zoom-in on a dendrite (B) and single synapses (C). (D-J) Quantification of Lphn1 nanoclusters by STED microscopy as shown in (B, C): mean number of Lphn1 nanoclusters per synapse (D); distribution of Lphn1 nanocluster numbers among synapses (E); nanocluster sizes (F), pre- and postsynapse areas (G, I) and vGLUT1- and Homer1-staining intensities normalized to ΔCre (H, J). All summary graphs are means ± SEMs. Numbers in bars show number of synapses/cultures analyzed. Statistics were done using paired t-tests, but no significant differences were detected.

**Figure 3:**
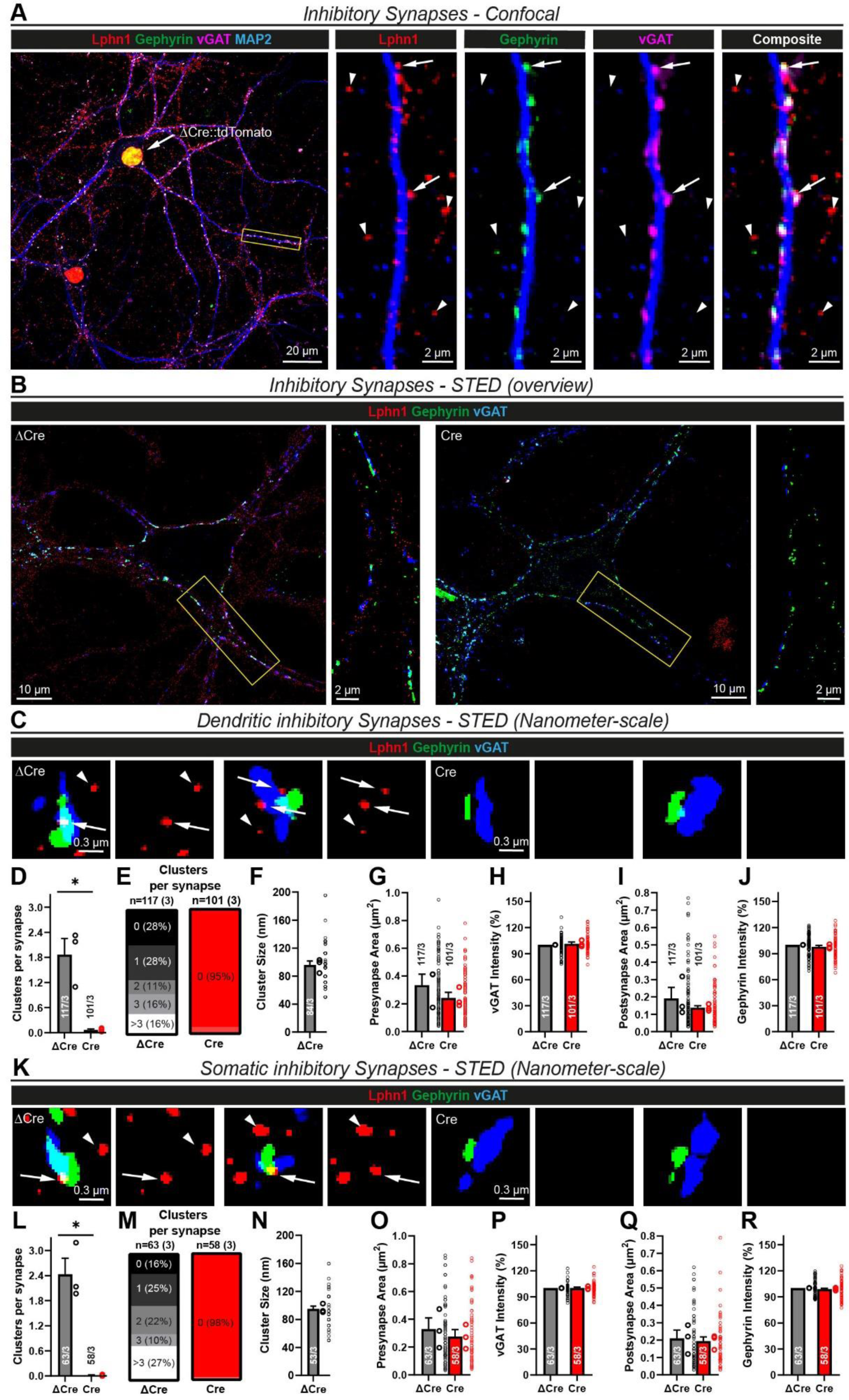
Lphn1 forms nanoclusters in dendritic and somatic inhibitory synapses similar to excitatory synapses. (A) Co-localization of myc-Lphn1 with inhibitory synapse markers using standard diffraction-limited confocal microscopy. After surface labeling for myc-Lphn1, Lphn1 cKO hippocampal cultures were permeabilized and stained for presynaptic (vGAT) and postsynaptic (Gephyrin) markers. Lphn1 is expressed in inhibitory synapses (arrows) but also present in puncta that do not stain for inhibitory synapse markers and could represent excitatory synapses (arrow heads). (B) STED microscopy of Lphn1 cKO hippocampal cultures reveals nanoclusters of Lphn1 in inhibitory synapses. Cultures were infected with lentiviruses encoding Cre recombinase or functionally deficient ΔCre recombinase, surface labelled for myc-Lphn1, then permeabilized and stained for presynaptic (vGAT) and postsynaptic (Gephyrin) markers of inhibitory synapses. Almost no nanoclusters can be found on neurons expressing Cre recombinase. (C) Single dendritic synapses from (B) shown at a smaller scale. Subsets of Lphn1 nanoclusters co-localize with inhibitory synapse markers (arrows) while others do not and are possibly from excitatory synapses (arrowheads). (D-J) Quantification of Lphn1 nanoclusters in dendritic inhibitory synapses as shown in (B, C): Mean amount of Lphn1 nanoclusters per synapse in (D) and number of synapses with no or a certain number of nanoclusters in (E), cluster diameter in (F), pre- and postsynapse areas in (G, I) and staining intensities of pre- and postsynaptic markers normalized to ΔCre in (H, J). (K) Single somatic synapses from (B) shown at a smaller scale. Lphn1 nanoclusters co-localize with inhibitory synapse markers (arrows) while others are possibly extrasynaptic (arrowheads). (L-R) Quantification of Lphn1 nanoclusters in somatic inhibitory synapses as shown in (B, K): Mean amount of Lphn1 nanoclusters per synapse in (L) and number of synapses with no or a certain number of nanoclusters in (M), Cluster Diameter in (N), pre- and postsynapse areas in (O, Q) and staining intensities of pre- and postsynaptic markers normalized to ΔCre in (P, R). All summary graphs (D-J and L-R) are means ± SEMs. Numbers in bars show number of synapses/cultures analyzed. Statistics were done using paired t-tests (*, p < 0.05).

Since conventional confocal microscopy is diffraction-limited and poorly resolves individual synapses, we aimed to confirm the presence of Lphn1 in both excitatory and inhibitory synapses using STED microscopy. To ensure that the Lphn1 signal we observe is specific, we examined both ΔCre (Lphn1-expressing) and Cre (Lphn1-deleted) hippocampal cultures and stained them as described above. Intriguingly, we found that Lphn1 forms nanoclusters in excitatory synapses (Figure 2B, C) as previously shown for other synaptic adhesion proteins, for example for the Lphn ligand Teneurin-3 (Zhang et al. 2022). The majority of synapses in the ΔCre condition displayed one to three nanoclusters, while the nanoclusters were absent in ∼90% of synapses in the Cre condition (Figure 2D, E). The nanoclusters had a mean diameter of ∼90-100 nm (Figure 2F), similar to the previously published dimensions of teneurin nanoclusters (Zhang et al. 2022). The deletion of Lphn1 did not significantly influence the area or staining intensity of pre- or postsynaptic specializations (Figure 2G-J), consistent with largely intact excitatory synapses after the Lphn1 deletion.

We next investigated inhibitory synapses. As suggested by conventional confocal microscopy, Lphn1 nanoclusters were prominently observed in inhibitory synapses, both on dendrites (Figure 3B, C) and the cell soma (Figure 3B, K). As a negative control, almost no clusters were found in inhibitory synapses in Cre-expressing neurons lacking Lphn1 (Figure 3D, L). Interestingly, in the ΔCre condition, the number of nanoclusters per synapse was on average higher in somatic (mean = ∼2.4 nanoclusters, Figure 3L, M) than in dendritic inhibitory synapses (mean = ∼1.8 nanoclusters, Figure 3D, E), which in turn was higher than the average number of nanoclusters in excitatory synapses (mean = ∼1.3 nanoclusters, Figure 2D, E). However, the nanocluster size of ∼90-100 nm was similar in all synapse types (Figure 2F, 3F, 3N). Pre- and postsynaptic areas had a small, statistically insignificant trend to being smaller in the Cre condition while the synapse-marker staining intensities were unchanged between ΔCre and Cre expressing neurons (Figure 3G-J and 3O-R).

These data suggest that Lphn1 is localized to the majority of both excitatory and inhibitory synapses in cultured hippocampal neurons and that it assembles into similar nanoclusters in these synapses, although it is on average more abundant in inhibitory synapses.

### Lphn1 is not required for neuronal morphogenesis

Earlier analyses of constitutive Lphn1 KO neurons suggested that Lphn1 interactions with teneurins promotes axonal outgrowth (Vysokov et al. 2018), whereas Lphn3 was proposed to mediate an axon-repellant signal (Pederick et al. 2021). Since functions in dendritic or axonal outgrowth could confound analysis of a role for Lphn1 in synapse assembly, we examined neuronal morphology in developing hippocampal neurons cultured from Lphn1 cKO mice. We lentivirally expressed Cre or ΔCre in the neurons and additionally sparsely transfected them with an EGFP-expression plasmid, which allowed us to quantify the size and shape of axons and dendrites as a function of the Lphn1 deletion (Figure 4A-G). However, we detected no significant morphological changes caused by the Lphn1 deletion. In particular, the total axon length (Figure 4B), number of axon branch points (Figure 4C), total dendrite length (Figure 4D), number of dendrite branch points (Figure 4E) and soma size (Figure 4F) were not altered in Lphn1-deficient neurons compared to control neurons analyzed in parallel. The GFP fluorescence intensity as a measure of transfection efficiency was also unchanged between groups (Figure 4G). Thus, in culture, the Lphn1 deletion had no effect on axonal or dendritic development.

**Figure 4:**
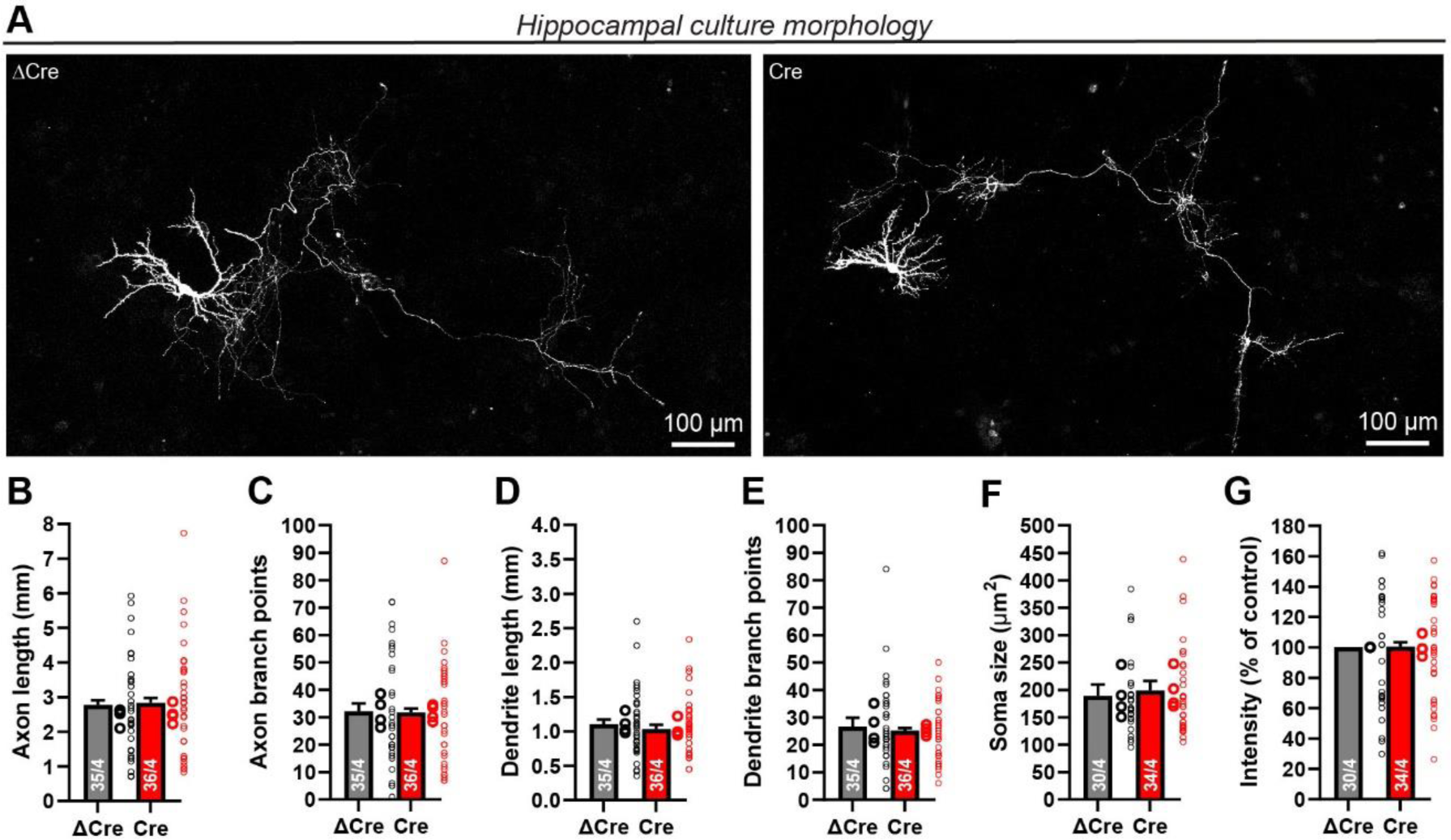
Lphn1 is not required for dendritic or axonal arborization. (A) Sample images of sparsely labeled cultured hippocampal neurons that had been infected with lentiviruses encoding active (Cre) or mutant inactive (ΔCre) Cre-recombinase. Sparse transfection of a CMV-promoter driven GFP allows tracing of dendrites and axons of individual neurons. Co-staining with MAP2 (not shown) was used to distinguish axons (MAP2 negative) and dendrites (MAP2 positive). White false color was used for the GFP signal for better contrast. (B-G) The Lphn1 deletion does not cause statistically significant changes in total axon length (B), axon branch points (C), total dendrite length (D) dendrite branch points (E) or soma size (F). Soma intensity was measured to show in G that similar amounts of GFP are expressed in both conditions and is displayed normalized to the ΔCre condition. All summary graphs (B-G) are means ± SEMs. Numbers in bars show number of neurons/cultures analyzed. Statistics were done using paired t-tests; no significant differences were noted.

### Lphn1 is also not required for excitatory synapse assembly

Previously, deletions of Lphn2 and Lphn3 were shown to decrease spine density and excitatory synapse numbers in hippocampal neurons both in culture and in vivo (Anderson et al. 2017, Sando et al. 2019). We therefore imaged EGFP-transfected cultured hippocampal neurons expressing or lacking Lphn1 at high resolution to monitor single spines (Figure 5A). Quantifications revealed only a minor trend towards a decreased spine density in Lphn1-deficient neurons, suggesting that different from the Lphn2 and Lphn3 deletions, the Lphn1 deletion does not significantly lower spine numbers (Figure 5B).

**Figure 5:**
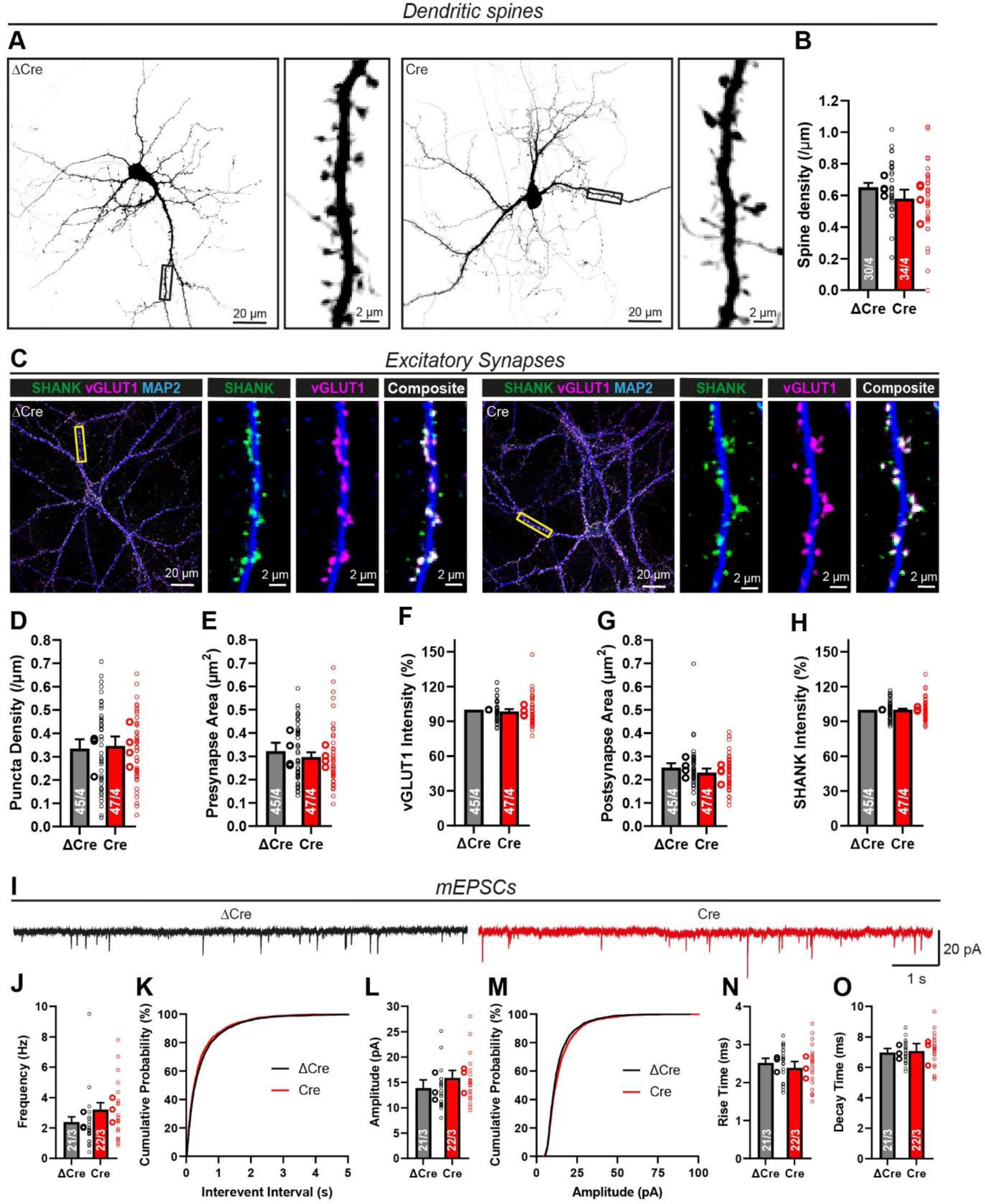
Lphn1 is not required for excitatory synapse formation or spine development. (A, B) Spine density analysis in cultured neurons. GFP transfected neurons have been imaged at sufficient resolution to count individual spines (A). Black false color was used for the GFP signal for better contrast. Only a minor decrease in spine density per µm of dendrite was observed, which was not statistically significant (B). (C) Immunocytochemistry of excitatory synapses in Lphn1 deficient cultures. Lphn1 cKO cultures were infected with lentiviruses encoding Cre recombinase or functionally deficient ΔCre recombinase, permeabilized and stained for presynaptic (vGLUT1) and postsynaptic (SHANK) markers of excitatory synapses as well as MAP2 to visualize dendrites. (D-H) Quantification of images as shown in (C). No significant changes were observed in any of the following parameters: (D) The density of synaptic puncta, defined by co-localized pre- and postsynaptic puncta per µm of dendrite; (E, G) area of pre- and postsynaptic puncta; (F, H) intensity of the staining of vGLUT1 and SHANK normalized to ΔCre. (I-O) Recordings of miniature excitatory postsynaptic currents (mEPSCs) in the presence of tetrodotoxin (TTX) show no significant difference after Lphn1 depletion. Representative traces are shown in (I), mEPSC frequency and cumulative plot of the interevent interval in (J, K), mEPSC amplitude and cumulative plot of the amplitude distribution in (L, M) and mEPSC kinetics in (N, O), defined as times of the event to rise from 10% to 90% of its amplitude and conversely to decay from 90% to 10% of its amplitude. All summary graphs (B-F, H and J-O) are means ± SEMs. Numbers in bars show number of neurons/cultures analyzed. Statistics were done using paired t-tests or Mann-Whitney tests (for electrophysiological recordings), but no significant differences were noted.

The lack of a change in spine numbers in Lphn1-deficient neurons suggests that, surprisingly, the Lphn1 deletion may not affect excitatory synapse formation, different from the Lphn2 and Lphn3 deletions (Anderson et al. 2017, Sando et al. 2019). Contrary to this suggestion, however, a previous study showed that cultured hippocampal Lphn1-deficient neurons exhibited a reduction in the staining intensities for presynaptic markers, suggesting a loss of synapses, although no measurements of synapse numbers were performed (Vitobello et al. 2022). To test this question directly, we stained Lphn1-deficient and control neurons by immunocytochemistry for vGLUT1 and SHANK, which are markers for pre- and postsynaptic excitatory synapses (Figure 5C), and measured the synapse density. Consistent with the lack of a change in spine density, the density of synaptic puncta was identical between control and Lphn1-deficient neurons (Figure 5D). The area and staining intensities of pre-and postsynaptic regions were also unchanged (Figure 5E-H).

The morphological analyses suggest that the Lphn1 deletion does not affect excitatory synapse numbers in pyramidal neurons. To verify these findings, we performed whole-cell patch-clamp recordings in control (ΔCre) and Lphn1-deficient (Cre) neurons in the presence of tetrodotoxin (TTX), and measured spontaneous miniature excitatory postsynaptic currents (mEPSCs) (Figure 5I). The mEPSC frequency and amplitude of these events were slightly, but not significantly elevated by the Lphn1 deletion (Figure 5J-M), while their kinetics were unchanged (Figure 5N, O). Recording parameters and passive cell characteristics were similar in both conditions throughout all electrophysiological experiments (Suppl. Figure 2).

The absence of an excitatory synapse phenotype in Lphn1-deficient pyramidal neurons is surprising in view of the robust Lphn1 expression in these synapses, the fact that an excitatory synapse phenotype was reported for a constitutive Lphn1 KO mouse (Vitobello et al. 2022), and the large decreases observed in excitatory synapse and spine numbers in Lphn2- and Lphn3-deficient pyramidal neurons (Anderson et al. 2017, Sando et al. 2019). To independently confirm this observation, we generated a second Lphn1 cKO mouse using ES cells obtained from the EUCOMM consortium (Figure 6A). The initial mice generated in this approach express LacZ from the endogenous Lphn1 gene (*Adgrl1*), enabling an independent assessment of Lphn1 expression. LacZ staining confirmed that Lphn1 is broadly expressed throughout the brain (Figure 6B). We then crossed the EUCOMM mice with a germline Flp recombinase-expressing mouse to generate Lphn1 cKO mice that contain normal Lphn1 levels (Figure 6A). Infection of cultured cKO hippocampal neurons with ubiquitin promoter-driven Cre recombinase resulted in a > 97% deletion of the floxed exon compared to a ΔCre infected control (Figure 6C). We additionally crossed the cKO mice with a Cre-dependent tdTomato reporter mouse line (Ai14), and analyzed the spine density in the Lphn1 cKO/Ai14 mice as a function of the Lphn1 deletion in vivo. Specifically, we stereotactically infected CA1 pyramidal neurons in neonatal Lphn1 cKO mice with Cre-expressing lentiviruses and filled infected Lphn1-deficient and uninfected control cells with biocytin via the patch pipette (Figure 6D, E). Subsequent analyses of the spine density as a proxy for synapse density again failed to detect any change in Lphn1-deficient neurons compared to control neurons in the *S. oriens*, *radiatum*, or *lacunosum-moleculare* (Figure 6F-I). These findings provide further support for the unexpected conclusion that Lphn1 is not required in hippocampal pyramidal neurons for synapse formation.

**Figure 6:**
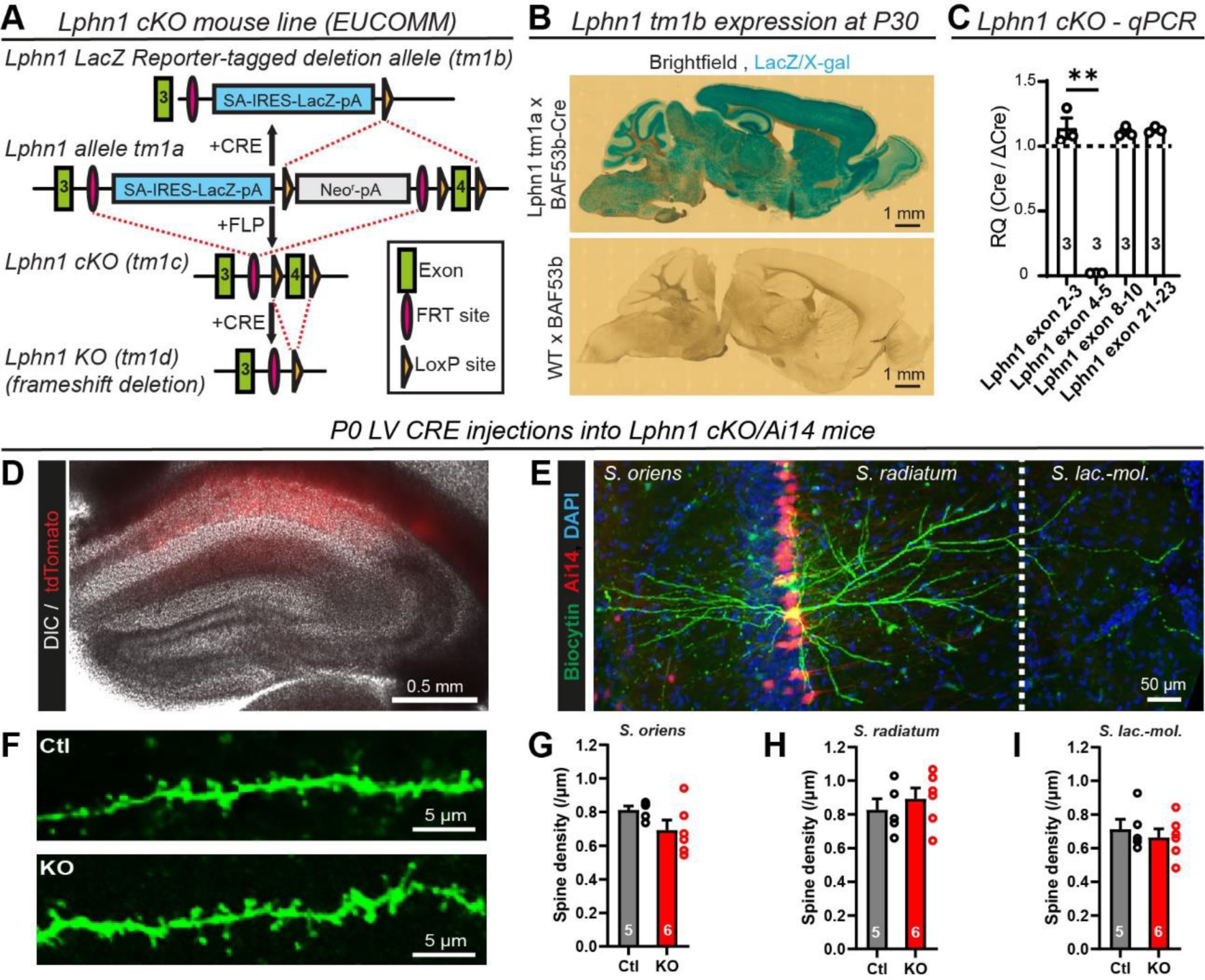
Sparse knockout of Lphn1 *in vivo* does not alter neuronal morphology or spine density. (A) Schematic design of Lphn1 cKO mice generated from EUCOMM ES cells (for details see methods). The original line (tm1a) allows for introduction of a LacZ reporter in the presence of Cre recombinase. Alternatively, a conditional allele (tm1c), in short Lphn1 cKO, can be generated with FLP recombinase followed by conditional Lphn1 deletion with Cre. (B) LacZ reporter staining in Lphn1 tm1a or wild-type mice expressing pan-neuronal BAF53B-Cre suggests high neuronal Lphn1 expression throughout the postnatal brain. (C) qPCR on mRNA samples from Lphn1 cKO primary hippocampal cultures infected with lentivirus encoding either Ubiquitin-driven GFP-deltaCre (control) or GFP-Cre confirms knockout of exon 4 in the Cre condition. (D-I) Morphological analysis of CA1 pyramidal cell dendrites infected with Cre recombinase lentivirus and non-infected controls in acute slices. (D) Overview image of a hippocampus, with Cre infected cells highlighted by Ai14 reporter expression in red. (E) A 20x image of a biocytin labelled CA1 pyramidal cell and all analyzed regions of interest; *Stratum oriens, Stratum radiatum, Stratum lacunosum moleculare*. (F) Representative image of dendritic spines in high magnification. (G-I) There was no significant difference in spine density in the *Stratum oriens* (G), *Stratum radiatum* (H), or *Stratum lacuosum moleculare* (I). All summary graphs are means ± SEMs. Numbers in bars show number of cultures (C) or mice (G-I) analyzed. Statistics were done using a paired t-test on culture values or Mann Whitney test on values of individual mice (**, p < 0.01).

### Lphn1 is essential for normal assembly of somatic inhibitory synapses

Since the expression of Lphn1 in inhibitory synapses seems to be higher than in excitatory synapses (Figure 2, 3), we next stained cultured Lphn1-deficient and control hippocampal neurons for the inhibitory synapse markers vGAT and Gephyrin (Figure 7A). Dendritic inhibitory synapses lacking Lphn1 were nearly unchanged in density, size or marker intensity (Figure 7B-F), with only a small decrease in size and intensity. When examining somatic inhibitory synapses (Figure 7G), however, we observed a dramatic decrease (∼50-60%) in puncta density (Figure 7H) as well as a minor decrease in pre- and postsynaptic area and staining intensity (Figure 7I-L).

**Figure 7:**
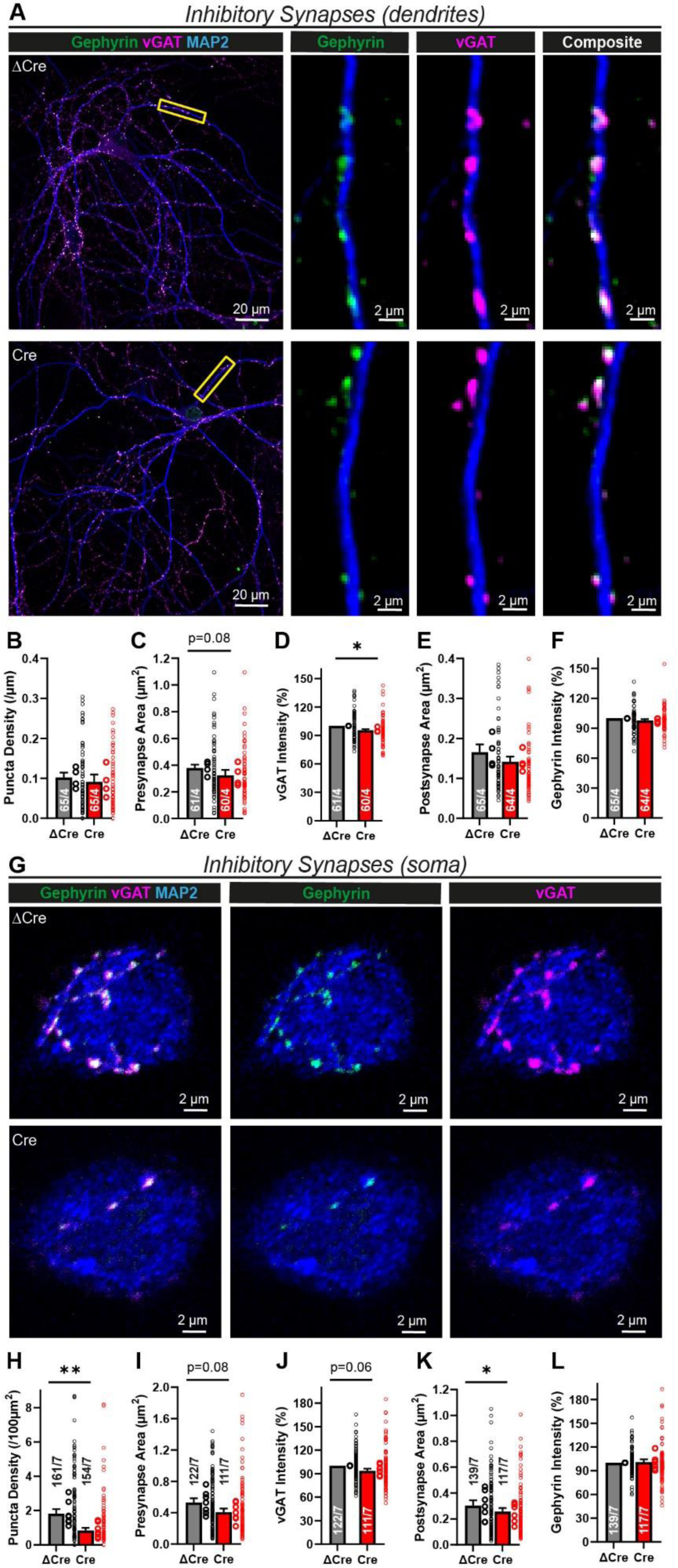
Lphn1 is specifically essential for somatic inhibitory synapse formation. (A) Immunocytochemistry of inhibitory synapses in Lphn1 deficient cultures. Lphn1 cKO cultures were infected with lentiviruses encoding Cre recombinase or functionally deficient ΔCre recombinase, permeabilized and stained for presynaptic (vGAT) and postsynaptic (Gephyrin) markers of inhibitory synapses as well as MAP2 to visualize dendrites. (B-F) Quantification of images as shown in (A). No major changes of the following parameters were observed in the Cre condition: (B) The density of synaptic puncta, defined by co-localized pre- and postsynaptic puncta per µm of dendrite; (C, E) area of pre- and postsynaptic puncta; (D, F) intensity of the staining of vGAT and Gephyrin normalized to ΔCre. (G) Stained cultures as described in (A) were additionally imaged at the cell soma, above the dendritic plane, to assess somatic inhibitory synapses. (H-L) Quantification of images as shown in (G). Somatic inhibitory synapse density, defined by co-localized pre- and postsynaptic puncta per µm^2^ of soma, was significantly reduced in the Cre condition (H), while the area of pre- and postsynaptic puncta as well as intensity of the staining of vGAT and Gephyrin (normalized to ΔCre) were only marginally reduced (I-L). All summary graphs (B-F and H-L) are means ± SEMs. Numbers in bars show number of neurons/cultures analyzed. Statistics were done using paired t-tests (*, p < 0.05; **, p < 0.01).

To ensure that the decrease in inhibitory innervation is not a secondary effect due to lower numbers of inhibitory neurons in cultures infected with Cre virus, we determined the percentage of GABAergic neurons using GAD67 staining (Suppl. Figure 3A). In both conditions we estimated an interneuron density of ∼10% (Suppl. Figure 3B). Staining intensity of GAD67 was also not significantly altered (Suppl. Figure 3C).

Thus, in contrast to Lphn2 and Lphn3, Lphn1 is not required for excitatory synapse formation but is essential for inhibitory synapse formation, in particular for somatic inhibitory synapses. To verify these findings, we performed patch-clamp whole-cell recordings of miniature inhibitory postsynaptic currents (mIPSCs) in control and Lphn1-deficient hippocampal neurons in the presence of TTX (Figure 8A). When we analyzed total mIPSCs, we observed a minor decrease in mIPSC frequency of ∼20% and a similar decrease in amplitude (Figure 8B-E), while the mIPSC kinetics were largely unchanged (Figure 8F, G). Interestingly, large amplitude mIPSCs seemed to be affected more severely than small amplitude mIPSCs (Figure 8A). mIPSCs originating from the cell soma are thought to exhibit larger amplitudes and faster rise times than mIPSCs generated in more distant dendrites (Wierenga and Wadman 1999). We therefore stratified mIPSCs into high-amplitude and low-amplitude events using an amplitude threshold, defined as the 50% percentile in the amplitude distribution of the ΔCre condition (average ∼32 pA). Intriguingly, the high-amplitude event frequency was reduced by ∼40-50% in Lphn1 KO neurons (Figure 8H, I), while low amplitude events were unchanged (Figure 8N, O). Amplitudes and kinetics of both subsets were not altered between the Cre and ΔCre conditions (Figure 8J-M and 8P-S).

**Figure 8:**
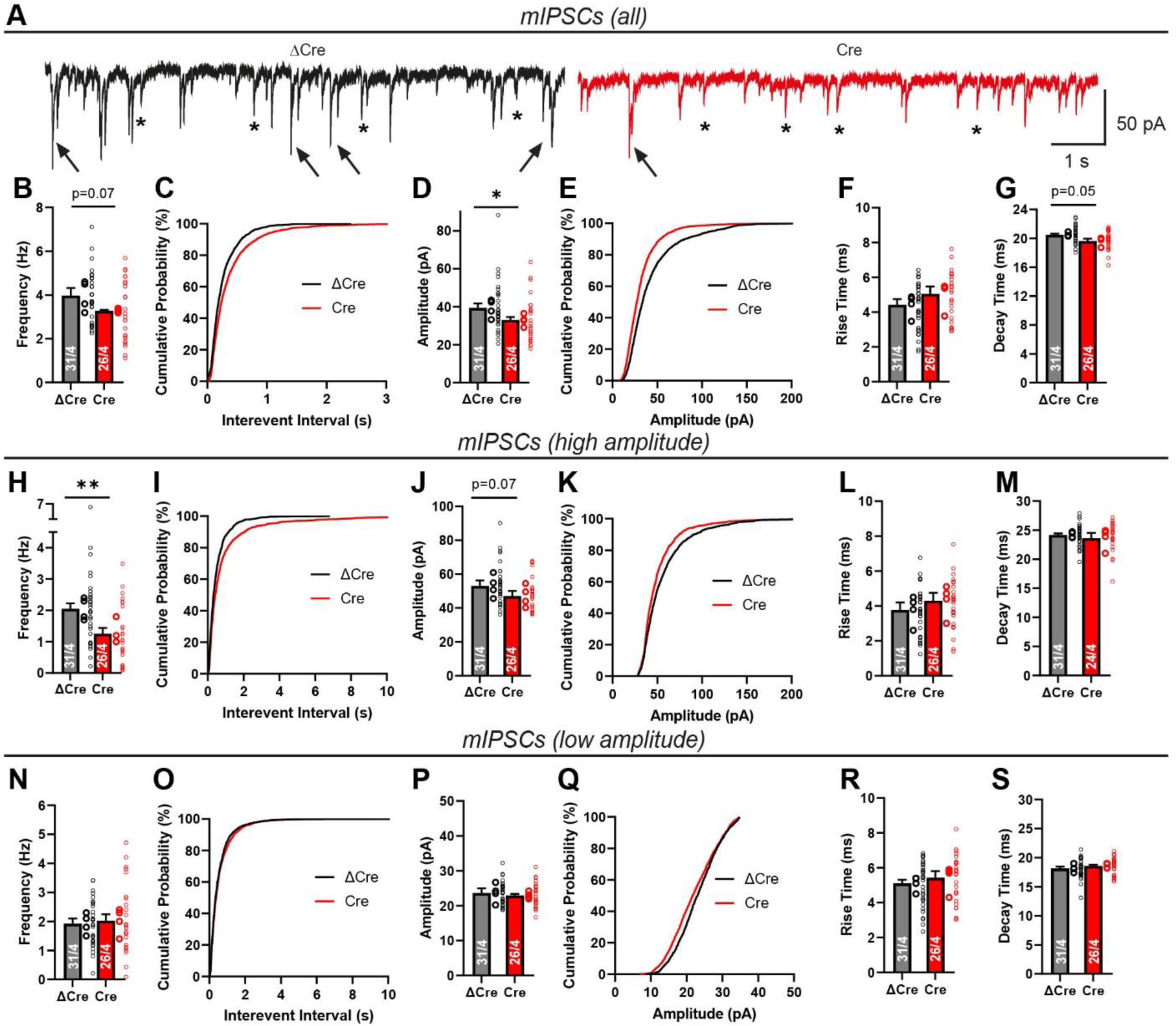
Electrophysiological analyses of cultured hippocampal neurons show that Lphn1 is essential for normal inhibitory synaptic transmission. Lphn1 cKO cultures were infected with lentiviruses encoding Cre recombinase or functionally deficient ΔCre recombinase and electrophysiological measurements were performed in whole-cell voltage clamp configuration: (A-G) Recordings of miniature inhibitory postsynaptic currents (mIPSCs) in the presence of tetrodotoxin (TTX) show reduced frequency and amplitude after Lphn1 depletion. Representative traces (A) show a specific reduction of high-amplitude mIPSCs (arrows), while low-amplitude mIPSCs are largely unchanged (asterisks). (B, C) mIPSC frequency is reduced in the Cre condition, and the cumulative plot of interevent intervals of Cre is shifted towards higher intervals. (D, E) mIPSC amplitude is significantly reduced and the cumulative plot of the amplitude distribution is shifted toward smaller amplitudes in the Cre condition. (F, G) mIPSC kinetics, defined as times of the event to rise from 10% to 90% of its amplitude and conversely to decay from 90% to 10% of its amplitude, show slightly higher rise times and smaller decay times. (H-S) Stratified analysis of high- and low-amplitude mIPSC. For each experiment, an amplitude threshold based on the 50% value of the control condition (ΔCre) was defined (∼32 pA on average) and applied to all analyzed events in both conditions. (H-M) Analysis of high-amplitude mIPSCs reveal a strong reduction in frequency after Lphn1 knockout (H, I), but no significant changes in amplitude, rise time and decay time (J-M). Low-amplitude events do not show any significant differences in frequency, amplitude or event kinetics (N-S). All summary graphs are means ± SEMs. Numbers in bars show number of neurons/cultures analyzed. Statistics were done using Mann-Whitney tests on individual neuron recordings (*, p < 0.05; **, p < 0.01).

To further validate an inhibitory synapse phenotype induced by the Lphn1 deletion, we next analyzed evoked inhibitory postsynaptic currents (eIPSCs) (Figure 9A). We observed a ∼50% decrease in eIPSC amplitude as well as a modest increase in the coefficient of variation (Figure 9B, C). These results indicate that the Lphn1 deletion strongly impaired inhibitory synaptic transmission. The effect of the Lphn1 deletion is probably more robust as measured by eIPSCs than as measured by mIPSCs because somatic inhibitory synapses are functionally more potent than dendritic inhibitory synapses since they are closer to the recording electrode. Rise and decay times of eIPSCs were not significantly altered between Cre and ΔCre expressing neurons (Figure 9D, E).

**Figure 9:**
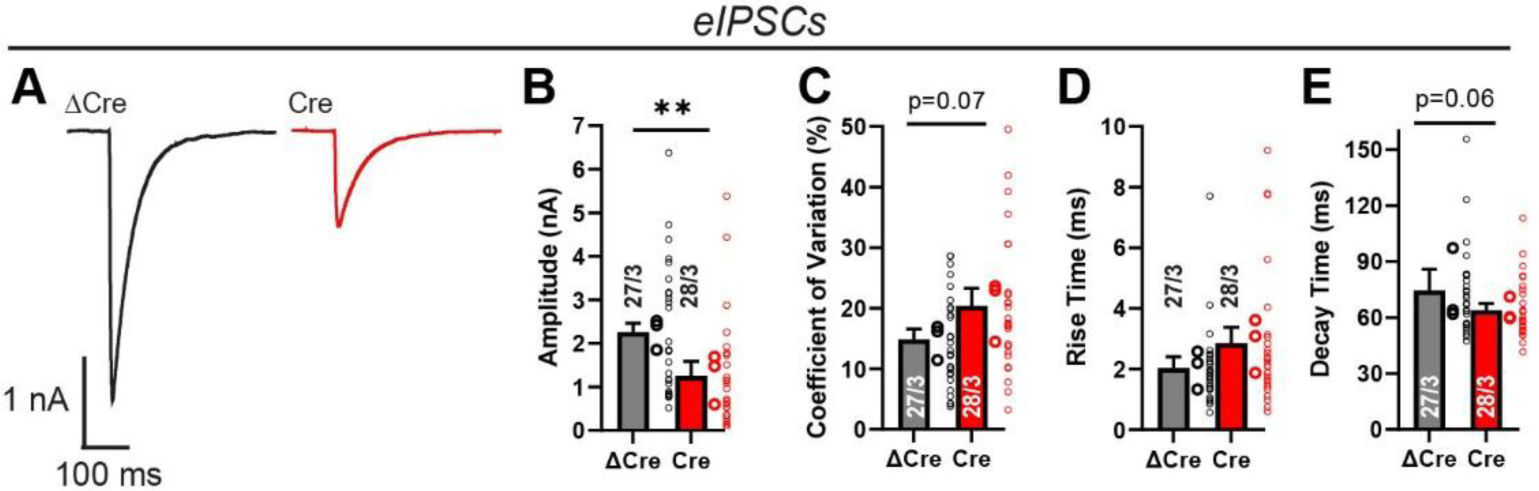
Electrophysiological measurements of evoked responses confirm essential role of Lphn1 in inhibitory synapses. Recordings of evoked inhibitory postsynaptic currents (eIPSCs) in Lphn1 knockout neurons show impaired inhibitory synaptic transmission. Responses of neurons to a 1 ms square pulse stimulus, delivered by a bipolar electrode to the culture coverslip in 100 µm distance from the cell body were recorded, while excitatory responses were pharmacologically suppressed. Representative traces are shown in (A). In the Cre condition, eIPSC amplitude is significantly reduced (B) and the coefficient of variation is slightly elevated (C). eIPSC kinetics (D and E), defined as times of the event to rise from 20% to 80% of its amplitude and conversely to decay from 80% to 20% of its amplitude, were largely unchanged. B-E show means ± SEMs. Numbers in bars show number of neurons/cultures analyzed. Statistics were done using Mann-Whitney tests on individual neuron recordings (**, p < 0.01).

## DISCUSSION

In the current study we generated conditional Lphn1 knockout mice that carry a knocked-in N-terminal myc epitope tag to enable us to immunolocalize the Lphn1 protein (Figure 1A-E). Using immunocytochemistry, we show that in the hippocampal CA1 region, Lphn1 is uniformly present throughout the dendritic tree of pyramidal neurons (Figure 1F, G), different from Lphn2 and Lphn3 that are enriched in the *S. lacunosum-moleculare* or the *S. oriens* and *S. radiatum*, respectively (Anderson et al. 2017, Sando et al. 2019). Moreover, we find in cultured hippocampal neurons using STED super-resolution microscopy that, unexpectedly, Lphn1 forms nanoclusters in both excitatory and inhibitory synapses (Figure 2, 3). The majority of excitatory and inhibitory synapses have at least one nanocluster, with a third of excitatory synapses (Figure 2E) and approximately 44% of dendritic and 59% of somatic inhibitory synapses featuring more than one Lphn1 nanocluster (Figure 3E, M). Consistent with the enhanced presence of Lphn1 in somatic inhibitory synapses, we found that in cultured hippocampal neurons the deletion of Lphn1 caused a significant decrease in the number of somatic but not dendritic inhibitory synapses (Figure 7). In parallel, we observed an impairment in inhibitory synaptic strength that most likely was due to the decrease in inhibitory somatic synapses. Specifically, measurements of mIPSCs revealed that the frequency of high-amplitude mIPSCs but not of low-amplitude mIPSCs that probably reflect synaptic events close to or distant from the soma, respectively, was significantly decreased (Figure 8H, N). Furthermore, the amplitude of evoked IPSCs was lower in Lphn1-deficient than control neurons without a significant change in the coefficient of variation (Figure 9). However, we observed no other major changes in Lphn1-deficient neurons. Neither dendritic nor axonal arborizations were changed (Figure 4), spine numbers were not decreased, excitatory synapse numbers were not lowered (Figure 5), and mEPSCs were not significantly changed (Figure 5). The lack of a change in spines was confirmed by in vivo deletion of Lphn1 using a second independent Lphn1 cKO mouse line, suggesting that it is not an artifact of one particular mouse line (Figure 6).

By demonstrating that Lphn1 is present in both excitatory and inhibitory synapses, Lphn1 emerges as one among few postsynaptic adhesion molecules that are in both classes of synapses, which otherwise share few postsynaptic components since their receptors and scaffolding molecules are different. With a diameter of approximately 90 nm, the sizes of the nanoclusters formed by Lphn1 in excitatory and inhibitory synapses are comparable (Figure 2F, 3F, 3N), suggesting a similar molecular organization. The Lphn1 nanoclusters we describe here using STED super-resolution microscopy closely resemble those previously observed using dSTORM for presynaptic teneurin-3 (Zhang et al., 2022), neurexins (Lloyd et al., 2023; Trotter et al., 2019), and neuroligins (Haas et al., 2018; Nozawa et al., 2022; Han et al., 2022), suggesting that such nanoclusters are a general feature of synaptic adhesion molecules. The detection of synaptic Lphn1 nanoclusters in both excitatory and inhibitory synapses also resolves the previously puzzling observation that teneurins, which are presynaptic latrophilin ligands, are broadly expressed in both excitatory and inhibitory neurons and could thus represent presynaptic interaction partners for postsynaptic Lphn1 in both types of synapses.

Our findings also raise important questions. First, are Lphn2 and Lphn3 also present in inhibitory synapses like Lphn1? This seems likely but has not yet been studied. Second and more puzzlingly, why doesn’t the Lphn1 deletion in our experiments produce a major phenotype in excitatory synapses, given that the Lphn2 and Lphn3 deletions cause major impairments in excitatory synapses and that previous studies on constitutive Lphn1 KO mice characterized significant impairments in excitatory synapse function (Vitobello et al. 2022)? The most plausible explanation for this discrepancy is that the Lphn1 deletions were analyzed on very different genetic backgrounds in mice and that analyses were performed on conditional vs. constitutive deletions. It seems unlikely that the discrepancy is due to a technical issue in our study or that of Vitobello et al. (2022) since we have established that our deletion indeed abolishes Lphn1 protein expression as expected from the genetic strategy (Figure 1A-E) and the phenotypes we observed in inhibitory synapses (Figure 7-9) and Vitobello et al. (2022) described in excitatory/inhibitory synapses appear to be very robust. Note, however, that another previous paper on constitutive Lphn1 knockout mice using neurotransmitter release from synaptosomes as an assay also did not detect a major change in excitatory synapses, although a synaptic density phenotype would have been missed in that study (Tobaben et al., 2002).

Another difference between the present results and another previous analysis of Lphn1 constitutive KO mice is that we did not detect changes in axonal outgrowth (Figure 4), whereas the constitutive Lphn1 KO neurons were found to display axonal growth defects (Vysokov et al. 2018). Such defects are interesting since functions in neuronal morphogenesis were also reported for the *C. elegans* latrophilin homolog LAT-1 (Matúš et al. 2022). However, only fluorescence intensities of stained axons in fields of view were measured in the previous analysis and axon lengths were not tracked in cultured neurons (Vysokov et al. 2018). While these methods can give hints towards neuronal functions, they are easily confounded by variations in culture density or composition. Moreover, it is puzzling that some experiments assign an attractive function of the latrophilin-teneurin interaction in axonal outgrowth (Vysokov et al. 2018), whereas others propose a repellent function (Pederick et al. 2021). Moreover, latrophilins were proposed to have an essential role in neuronal migration during brain development (Del Toro et al. 2020), even though major developmental effects on cortical layers by the constitutive Lphn1 deletion were not apparent (Tobaben et al. 2002; Vysokov et al. 2018; Vitobello et al. 2022). At present these contradictions cannot be resolved although – again – genetic backgrounds may play a role, an issue that is challenging to resolve.

In summary, our results reveal the presence of similar Lphn1 nanoclusters in both excitatory and inhibitory synapses, with most synapses in hippocampal cultures featuring at least one nanocluster. Moreover, our findings uncover an essential function for Lphn1 in inhibitory synapses. These results expand our view of the synaptic role of latrophilins, suggesting a general function in most synapses independent of their transmitter type. However, major questions arose that need to be addressed in future, such as that of the role of Lphn1 in excitatory synapses. Such a role might be occluded by redundancy, or Lphn1 might be without function in these synapses. Given the differences in results with distinct mouse lines and the possibility of major genetic background effects, a more reductionist approach may be needed to deconstruct the contribution of latrophilins in general and Lphn1 in particular to different types of synapses. For example, it is possible that latrophilins function in the same pathway as other aGPCRs, such as BAI’s and CELSRs that also have synaptic functions (Najarro et al. 2012, Sigoillot et al. 2015, Wang et al. 2020, Wang et al. 2021, Bolliger et al. 2011, Shiu et al. 2022, Tu et al. 2018, Zhu et al. 2015, Stephenson et al. 2013, Aimi et al. 2023, Martinelli et al. 2016, Kakegawa et al. 2015, Freitas et al. 2023, Li et al. 2022, Zhou et al. 2021, Thakar et al. 2017), and that a more extensive deletion of multiple genes will be necessary to unravel the making of a synapse!

## MATERIALS AND METHODS

### Animal procedures

All mice housed at Stanford were weaned at 20-21 days of age and housed in groups of 2 to 5 on a 12 hr light/dark cycle with access to food and water ad libidum. All procedures conformed to National Institutes of Health Guidelines for the Care and Use of Laboratory Mice and were approved by the Stanford Animal Use Committees [Administrative Panel for Laboratory Animal Care (APLAC) / Institutional Animal Care and Use Committee (IACUC)]. Mice housed at Vanderbilt were weaned at 18-21 days of age and housed in groups of 2 to 5 on a 12 h light/dark cycle with food and water ad libidum. Vanderbilt Animal Housing Facility: All procedures conformed to National Institutes of Health Guidelines for the Care and Use of Laboratory Mice and were approved by the Vanderbilt University Administrative Panel on Laboratory Animal Care. Ai14 mice were obtained from Jackson laboratories (#7914) and were crossed to L1f cKO mice to obtain homozygous L1f cKO/homozygous Ai14 reporter. Baf53b-Cre mice (Actl6b-Cre) were obtained from Jackson laboratories (#27826). Mice were bred on a hybrid background to avoid penetrance of background mutations in inbred mouse strains.

### Generation of Lphn1 conditional knockout mice

Lphn1 cKO mice were generated from EUCOMM ES cells (Adgrl1^tm2a(EUCOMM)Hmgu^, parental cell line JM8A3.N1 (B6N)) at the Janelia Farm Research Campus. Founder mice were crossed to FLP mice (B6N.Cg-Tg(ACTFLPe)9205Dym/CjDswJ #19100) to remove the selection cassette and LacZ reporter, generating a conditional floxed exon 4. Lphn1 cKO/FLP mice were crossed to C57/Bl6J (Jackson #664) to remove the FLP allele, and heterozygous Lphn1 cKO non-littermates subsequently crossed to homozygosity. Lphn1 cKO mice are available via Jackson (#035185). The following primers were used to genotype Lphn1 cKO alleles:

Reaction 1: 5’-AGGCATCCTTATCCATGGAG-3’, 5’-TGCTATGGAGTGCAGAGACT-3’, 5’-CTCCTACATAGTTGGCAGTG-3’ (WT 374 bps, cKO 259bps)
Reaction 2: 5’-CTTGATCCAGTACACCTGTG-3’, 5’-GGCTCTGAAGGACTTTAGCA-3’ (WT 237 bps, cKO 317 bps)

Myc-tagged Lphn1 cKO mice were custom generated at Janelia Farm Research Campus. In brief, Exon 2 of Lphn1 was duplicated, with one duplicated exon 2 containing a myc tag inserted after the signal peptide, and the second exon 2 containing a myc-SNAP tag inserted following the signal peptide. Preferred frt sites flank the Neomycin selection cassette and Exon 2 containing myc-SNAP tag. Non-preferred frt3 sites flank the Exon 2 containing myc tag and Neomycin cassette. Both duplicated exon 2 are flanked by loxP1 sites to allow conditional deletion. Founder mice were crossed to B6N.Cg-Tg(ACTFLPe)9205Dym/CjDswJ #19100 and a PCR-based strategy was used to discriminate between FLP-mediated recombination events to select for myc-Exon2 containing Lphn1 mice. myc-Lphn1 mice identified as positive via PCR were subsequently crossed to C57/Bl6J to remove the FLP allele. Myc-Lphn1 cKO mice are available via Jackson (#035181). The following primers were used for genotyping the myc-Lphn1 allele:

Reaction 1: 5’- CATAGATgttaacGGATCCACC-3’, 5’- CCGGTACTGTAAGCTTTGTAG-3’ (mutant 222 bps)
Reaction 2: 5’- GACACACAGTTGTGACTGAC-3’, 5’- GCATCGCAGATCTTGTCATC-3’ (WT 359 bp, mutant 527 bp)
Reaction 3: 5’- GCACGTCACACTGACATTGT-3’, 5’- TCTGTGAGCTCCTACCTGAA-3’ (WT 271 bp, mutant 367 bp)

### Antibodies

Concentrations of antibodies that have been used for different experimental approaches in this study (immunohistochemistry = IHC; immunocytochemistry = ICC; immunoblot = IB) were: Avidin-Alexa Fluor 488 (Invitrogen, #S32354) 1:1000 (IHC), rabbit anti-c-myc (Sigma) 1:500 (ICC, IHC), 1:1000 (IB); mouse anti-beta-actin (Sigma) 1:5000 (IB), chicken anti-MAP2 (Encor) 1:2000 (ICC), guinea pig anti-vGLUT1 (SySy) 1:1000 (ICC), mouse anti-pan-SHANK (Neuromab) 1:1000 (ICC), guinea pig anti-vGAT (SySy) 1:1000 (ICC), mouse anti-Gephyrin (Neuromab) 1:1000 (ICC), mouse anti-Homer1 (SySy) 1:1000 (ICC), rabbit anti-NeuN (Millipore) 1:1000 (ICC), mouse anti-GAD67 (Millipore) 1:1000 (ICC), donkey anti-rabbit IRDye800CW (Licor, Cat# 926-32213) 1:10000 (IB), donkey anti-mouse IRDye680LT (Licor, Cat# 926-68022) 1:10000 (IB), goat anti-chicken AlexaFluor405 (ThermoFisher, Cat# A48260) 1:1000 (ICC), goat anti-chicken AlexaFluor546 (ThermoFisher, Cat# A11040) 1:1000 (ICC), goat anti-rabbit AlexaFluor405 (ThermoFisher, Cat# A31556) 1:1000 (ICC), goat anti-rabbit AlexaFluor546 (ThermoFisher, Cat# A11035) 1:1000 (ICC), goat anti-mouse AlexaFluor488 (ThermoFisher, Cat# A11029) 1:1000 (ICC), goat anti-mouse AlexaFluor647 (ThermoFisher, Cat# A21236) 1:1000 (ICC), goat anti-guinea pig AlexaFluor647 (ThermoFisher, Cat# A21450) 1:1000 (ICC), goat anti-rabbit STAR RED (Abberior, Cat# STRED-1002) 1:250 (ICC), goat anti-guinea pig STAR ORANGE (Abberior, Cat# STORANGE-1006) 1:500 (ICC), goat anti-mouse STAR 460L (Abberior, Cat# ST460L-1001) 1:500 (ICC).

### Lentiviral production

HEK293T cells (ATCC) were grown in a 37°C and 5% CO_2_ atmosphere in DMEM (Gibco Cat# 11995065) + 10% FBS. Lentiviral constructs expressing NLS-GFP-ΔCre or NLS-GFP-Cre driven by the human Synapsin promoter were described previously (Kaeser et al. 2011). In this study, similar constructs from the Südhof lab have been used (NLS-tdTomato-ΔCre or NLS-tdTomato-Cre under the influence of the human Synapsin promoter or NLS-GFP-ΔCre or NLS-GFP-Cre under the influence of the Ubiquitin promoter). Lentiviral packaging was achieved by transfecting 10.8 μg of the respective shuttle vectors in addition to helper plasmids (3 μg of pRSV-REV, 7.35 μg of pMDLg/pRRE and 3.84 μg of vesicular stomatitis virus G protein (VSVG)) into 75 cm^2^ of 40-50% confluent HEK293T culture using a calcium-phosphate approach. DNA was diluted in 450 μl of ddH2O and 50 μl of 2.5 M CaCl2 solution was added. Then the DNA-CaCl2 mix was added dropwise into an equal volume of 2xHEPES-buffered saline (274 mM NaCl, 10 mM KCl, 1.5 mM Na2HPO4, 12 mM Glucose, 42 mM HEPES, adjust pH to 7.05 using NaOH), while gently vortexing the tube. After 20 min of incubation at room temperature, the transfection mix was briefly flicked and added dropwise to the cells. After 24 h, medium was exchanged to neuron growth medium (components in section below) and after 48 h virus-containing medium was filtered through 0.22 μm PES membranes (GenClone, Cat# 25-243), aliquoted and stored at −80°C for application in culture experiments. For each virus batch the lowest necessary infectious titer was determined to infect > 95% of cultured neurons and corresponding quantities were added in all culture experiments. For lentiviral concentration, viral conditioned media was centrifuged at 5000 × g for 5 min to pellet cellular debris, filtered and ultracentrifuged at 55,000 × g for 1.5 hr through a sucrose cushion. Pellets were resuspended in MEM at 1/500 of the initial volume, aliquoted and stored at −80°C.

### Primary hippocampal cultures

P0 hippocampus dissection was performed in ice-cold HBSS (Hanks Balanced Salts (Sigma Cat# H2387-10X1L), 1 mM HEPES (pH 7.3), 4 mM NaHCO3), and tissue was digested in papain solution (100 μl papain suspension (Worthington, Cat#LS003127) in 5 ml HBSS) for 30 mins at 37 °C. Tissue was then washed 2 × in HBSS, 1 × in plating medium (5% fetal bovine serum (Sigma, Cat# F0926), B27 supplement (Gibco Cat#17504044), 0.4% glucose and 2 mM glutamine diluted in 1x MEM (Gibco Cat#51200038)) and thoroughly triturated. After filtration through a 70 μm cell strainer (Falcon Cat#352350), cells were suspended in plating medium either in 6-well plates coated with 0.05 M poly-D-lysine (Sigma, Cat# P7280) diluted in 0.05 M borate buffer (for protein extraction) or in 24-well plates containing 0 thickness glass coverslips (Assistant Cat#01105209 coated with Matrigel (Corning Cat#356235). On DIV1 80% of medium was exchanged to neuron growth medium (5% fetal bovine serum (Sigma, Cat# F0926), B27 supplement (Gibco Cat#17504044) and GlutaMAX™ (Gibco Cat#35050061) in Neurobasal A (Gibco Cat#10888022)). On DIV3 and DIV8, 50% of medium was exchanged for neuron growth medium containing 4 μM cytosine arabinofuranoside (Santa Cruz Biotechnology Cat#221454A). Lentivirus infections were performed during the media change on DIV3 (if not stated otherwise) and sparse calcium-phosphate transfections on DIV8. Neuronal cultures were kept in a 37°C and 5% CO2 atmosphere and analyzed on DIV14-16.

### Protein collection, SDS-PAGE and Immunoblotting

To extract total protein, neuronal cultures were washed twice with ice-cold PBS, and lysed for 20 min with freshly prepared RIPA buffer (50 mM Tris-HCl (pH 8), 1 mM EDTA, 150 mM NaCl, 0.5% Sodium deoxycholate, 0.1% SDS, 1% TritonX, PMSF 1 mM and cOmplete protease inhibitor cocktail (Sigma Cat# 11873580001)) on ice, under light agitation. The samples were then spun down in a tabletop centrifuge (precooled to 4°C) at 15000 g for 20 min and pellets were discarded. Concentrations of protein samples were measured with the Micro BCA™ Protein Assay Kit (Life Technologies Cat#23235), using a BSA standard curve. Equal amounts of protein were diluted in 5x sample buffer (50 mM Tris-HCl pH 6.8, 10% SDS, 40% glycerol, 100 mM DTT and bromophenol blue), loaded on 4–20% Mini-PROTEAN® TGX™ Precast Gels (BioRad) together with Xpert 2 Prestained Protein marker (GenDEPOT, Cat# P8503) and run at 25 mA/gel constant current. The Trans-Blot Turbo Transfer System and Buffer (BioRad, Cat# 1704150) was used following manufacturer’s instructions to perform a semi-wet transfer onto a 0.45 µm Amersham™ Protran® nitrocellulose membrane (Sigma, Cat# GE10600002). For blocking, membranes were incubated for 1 hr at 22°C in 5% non-fat dry milk/TBS (20 mM Tris-HCl pH 7.4, 150 mM NaCl). Membranes were then incubated overnight at 4°C in primary antibody diluted in 5% milk/TBST (TBS + 0.05% Tween), washed 3 × 5 mins with TBST, incubated for 45 min at 22°C with corresponding secondary antibodies diluted in TBST containing 0.01% SDS, washed 5 × 5 mins with TBST, and imaged using a Licor Odyssey system.

### Quantitative RT-PCR

Primary hippocampal cultures were generated from postnatal day 0 Lphn1 cKO (EUCOMM) mice as described above and infected at DIV1 with lentiviral Ubiquitin-driven GFP-CRE or GFP-ΔCRE. At DIV14, total mRNA was isolated using the Qiagen RNAeasy mini kit (#74104) and mRNA concentration and purity via A260/280 ratio measured using a Nanodrop (ThermoFisher). qPCR measurements were performed with VeriQuest Probe One-Step qRT-PCR Master Mix (Affymetrix #75700) and 10 ng mRNA template using an Applied Biosystems 7900HT apparatus using predesigned PrimeTime qPCR probe assays (IDT). The following probes were used; *Mm* Lphn1: Mm.PT.58.42048881 (exon 2-3); Mm.PT.58.6572453 (exon 4-5); Mm.PT.58.30545738 (exon 8-10); Mm.PT.58.10233084 (exon 21-23); *Mm* Gapdh: Mm.PT.39a.1. RQ was calculated via RQ =2^−[Ct(target, CRE)−Ct(target, ΔCRE)]−[Ct(Gapdh, CRE)−Ct(Gapdh, ΔCRE)]^.

### Sparse transfections of neuron cultures

In order to analyze neuronal morphology, neurons need to be visualized in their entirety without overlapping dendrites or axons. Therefore, sparse transfection using a calcium phosphate-based approach was performed to deliver a CMV promoter driven GFP plasmid into individual neurons. Per well in a 24 well plate, the following mixture was prepared: 0.5 μg of DNA, 1.5 μl of 2.5 M CaCl_2_ and ddH_2_O to a total volume of 15 μl. While gently vortexing a tube containing an equivalent volume of 2xBBS pH 7 (50 mM BES, 280 mM NaCl, 1.5 mM Na2HPO4), the DNA/calcium phosphate mixture was added dropwise and left to precipitate on room temperature for 15 min. 75% (750 μl) of neuron growth medium was taken off the cultured neurons and kept at 37°C. Neurons were then washed 3 × with prewarmed MEM (Gibco Cat#51200038) in the form of 75% media exchanges. After the final wash, 500 μl of MEM was kept in each well. The transfection mix was briefly vortexed before 30 μl were added dropwise per well, followed by 10 min of incubation at 37°C and 5% CO_2_. Thereafter, three 75% media exchanges for fresh prewarmed MEM were performed, leaving 100 μl per well after the last wash. 500 ul of saved preconditioned medium and 400 ul of fresh neuron growth medium was added to each well and the culture was put back into a 37°C and 5% CO_2_ atmosphere.

### Immunocytochemistry of cultured cells

For surface labeling, cells were washed 1 × in bath solution (see composition in section below) and incubated for 20 min at room temperature in primary antibody. After being washed 3 × with bath solution, neurons were then fixed in 4% PFA diluted in bath solution for 20 min at room temperature, washed 3 × in PBS, blocked for 60 min in 5% goat serum in PBS and stained with appropriate secondary antibody, diluted in 5% goat serum in PBS. Cells were then washed 4 × with PBS, fixed a second time in 4% PFA diluted in PBS for 5 min at room temperature and washed 3 × in PBS before permeabilization and subsequent staining. If no surface labeling was performed, cells were washed 1 × in bath solution, fixed for 20 min at room temperature in 4% PFA diluted in bath solution and washed 3 × in PBS. Cells were permeabilized/blocked for 60 min at room temperature in 0.2% TritonX100/5% goat serum/PBS and incubated in primary antibodies diluted in 0.2% TritonX100/5% goat serum/PBS overnight at 4 °C. The next day, neurons were washed 3 × with PBS and incubated in corresponding secondary antibodies in 0.2% TritonX100/5% goat serum/PBS for 1 hr at room temperature. After 4 × washing in PBS, coverslips with cells were briefly submerged into ddH2O to remove salts and mounted on Diamond White Glass microscope slides (Globe Scientific, Cat# 1358W) in a drop of Fluoromount-G (Southern Biotech Cat#0100-01) or ProLong™ Gold Antifade Mountant (ThermoFisher, Cat# P36930), if STED microscopy was to be performed. Slides were left to dry in the dark at room temperature for 48 hrs and then stored at 4 °C until imaging.

### Culture electrophysiology

Culture electrophysiology was performed essentially as previously described (Maximov et al. 2007) in whole-cell patch clamp configuration, holding the cell at −70 mV. Glass pipettes with a resistance of 2-4 MΩ were pulled from borosilicate glass capillaries (World Precision Instruments, Cat# TW150-4) using a PC-10 pipette puller (Narishige). A Multiclamp 700B amplifier and Digidata 1550A Low-Noise Data Acquisition Digitizer with Clampex 10 data acquisition software (Molecular Devices) were used to monitor synaptic currents, sampled at 4 kHz. For triggering evoked responses, focal square pulse stimuli of 1 ms duration were applied ∼100 μm from the cell soma via a bipolar electrode (FHC, Bowdoinham, ME) controlled by a Model 2100 Isolated Pulse Stimulator (A-M Systems, Inc.). Coverslips with neurons were put into extracellular bath solution that was composed of (in mM) 140 NaCl, 5 KCl, 2 MgCl2, 10 D-glucose, 10 HEPES (pH 7.4, adjusted with NaOH and 290-300 mOsm). For recording excitatory events 2 mM CaCl2 and for inhibitory events 0.5-0.75 mM CaCl2 was added. To pharmacologically isolate excitatory postsynaptic currents, the GABAA receptor blocker picrotoxin (50 μM) was added to the extracellular bath solution, while inhibitory postsynaptic currents were observed in the presence of AP5 (50 μM) and CNQX (10 μM) to block NMDA or AMPA receptors, respectively. To block action potentials during the recording of spontaneous miniature excitatory / inhibitory postsynaptic currents (mEPSCs / mIPSCs) 1 μM tetrodotoxin was added to the bath solution. Drugs mentioned above were obtained from Tocris (Minneapolis, MN, USA). The pipette solution for excitatory recordings contained (in mM) 135 Cs-Methanesulfonate, 8 NaCl, 10 HEPES, 0.3 EGTA, 0.3 Na2GTP, 2 MgATP, 7 phosphocreatine, 0.1 spermine, and 10 QX-314 (Tocris Cat# 1014) (pH 7.3, adjusted with CsOH and 306 Osm). For inhibitory recordings, the pipette solution was composed of (in mM) 135 CsCl, 10 HEPES, 1 EGTA, 4 MgATP, 0.4 Na2GTP, 7 phosphocreatine, 1 spermine and 10 QX-314 (Tocris Cat# 1014) (pH 7.3, adjusted with CsOH and 283 mOsm). Traces were analyzed while blinded to the experimental conditions using Clampfit 10 software (Molecular Devices). mEPSCs / mIPSCs were automatically detected in a template-based search and visually inspected for inclusion or rejection.

### Sparse lentiviral infections and neonatal injections

P0 Lphn1 cKO/Ai14 mutant mice were anesthetized for 5 min on ice. Concentrated lentivirus was injected with a glass pipette using an infusion pump (Harvard Apparatus). The hippocampal CA1 region was unilaterally targeted using the following coordinates from Lambda: anterior-posterior, +1.0 mm; medial-lateral, ±1.1 mm; and dorsal-ventral serial injections at, −1.5 mm, −1.3 mm, and −1.1 mm. Flow rate was 1μL/min and injected volume was 0.3 μl. Efficiency and localization of viral expression was confirmed by nuclear GFP expression under a fluorescent microscope.

### Patching acute brain slices

For achieving whole-cell patch clamp configuration, patch pipettes were pulled from borosilicate glass capillary tubes (TW150-4; World Precision Instruments) using a PC-10 pipette puller (Narishige). Pipette resistance with intracellular solution ranged from 3-5 MΩ. Lentivirus was injected into P0 mice. Infected CA1 pyramidal neurons were analyzed at P18-P30. Transverse hippocampal slices (300 μm) were prepared by cutting in ice-cold solution containing 228 mM sucrose, 2.5 mM KCl, 1 mM NaH_2_PO_4_, 2 6 mM NaHCO_3_, 0.5 mM CaCl_2_, 7 mM MgSO_4_, and 11 mM d-glucose saturated with 95% O_2_/5% CO_2_. Slices were then transferred to a holding chamber containing ACSF: 11mM NaCl, 2.5 mM KCl, 1 mM NaH_2_PO_4_, 26 mM NaHCO_3_, 2.5 mM CaCl_2_, 1.3 mM MgSO_4_-7H_2_O, 11 mM d-glucose, and ∼292 mOsm. Slices were recovered at 32°C for 30 minutes in a water bath, then at room temperature for at least 1 hour before recording. Slices were then transferred to a recording chamber where they were continuously perfused with oxygenated ACSF maintained at 32°C. Whole cell pipette solution was used containing 146 mM CsCl, 10 mM HEPES, 0.25 mM EGTA, 2 mM MgATP, 0.3 mM Na_2_GTP, 0.1 mM spermine, and 7 mM phoshocreatine. The pH of the solution was adjusted to 7.25-7.3 with 1M CsOH and osmolarity was between 294-298 mOsm.

### Immunohistochemistry of cryosections

During whole-cell patch clamp configuration, cells were filled for 10-15 minutes with 2 mg/mL biocytin (Sigma Aldrich #501787307) in whole patch solution. The recording pipette was slowly removed from the cell after recording was completed. Acute slices were washed briefly with PBS to remove excess ACSF solution then transferred to a 4% PFA (Electron Microscopy Science Cat# 15714) in PBS solution in 4°C overnight. Slices were then washed with PBS 5x for 5 min. Permeabilization solution (0.3% Triton X-100 in PBS) was applied for 30 minutes at room temperature. Slices were then placed in 5% normal goat serum (Jackson Immunoresearch #005000121) + 0.1% Triton X-100 for 1 hour at room temperature. Avidin-Alexa 488 (Invitrogen #S32354) (1:1000) in blocking solution was added to slices for incubation at room temperature for 1.5 hours. Slices were washed 1x for 5 min with DAPI (Sigma Cat# 10236276001;1:1,000) in PBS, 3x for 5 min with PBS, and then mounted on UltraClear microscope slides (Denville Scientific Cat# M1021) using 10 μL ProLong Gold antifade reagent (Invitrogen, #P36930).

### Imaging

Standard confocal imaging was performed on a Nikon A1 Eclipse Ti confocal microscope with 10x(air)/20x(air)/60x(oil) objectives (Apo, NA 1.4) and NIS-Elements AR acquisition software. Experiments were performed with constant laser intensities and acquisition settings for all conditions. Optimal Nyquist xy-resolution and Z-stacks with 0.3 μm / 1 μm spacing were used for visualizing synaptic puncta or spines / neuronal morphology, respectively. Confocal image analysis was mostly conducted using NIS Elements and ImageJ/Fiji. Synaptic puncta and spines on cultured neurons were counted on multiple secondary and tertiary dendrites and these values were averaged per neuron. Imaris software was used for automatic tracing of axons and dendrites. 2D-STED microscopy was performed on a Nikon Ti2-E microscope stand equipped with a CFI PLAN APO LAMBDA 100x(oil) objective (NA 1.45) and a STEDYCON confocal and STED module (Abberior). Pulsed excitation lasers were of 488 nm / 595 nm / 640 nm wavelength and a 775 nm pulsed STED laser was used for depletion, allowing for 60 nm resolution in all channels. Photons were detected using Time-gated Avalanche Photodiodes with 15-line accumulations and pixel size of 30×30 nm. Raw images were processed in Hugyen’s Essentials (SVI) using express deconvolution and exported as ICS files for further analysis in NIS Elements and ImageJ/Fiji. GAD67/NeuN stained neurons and Myc-stained brain slices were imaged using a VS120 slide scanner (Olympus) at 20x(air) magnification, exported as 8-bit OME files and analyzed in NIS Elements and ImageJ/Fiji. Dendritic spine images in brain slices were acquired using a Nikon A1r resonant scanning Eclipse Ti2 HD25 confocal microscope with 10x (Nikon #MRD00105, CFI60 Plan Apochromat Lambda, N.A. 0.45), 20x (Nikon #MRD00205, CFI60 Plan Apochromat Lambda, N.A. 0.75), and 60x (Nikon #MRD01605, CFI60 Plan Apochromat Lambda, N.A. 1.4) objectives, operated by NIS-Elements AR v4.5 acquisition software. For biocytin analysis, images were collected with the following resolution: 20x – 0.86 µM/pixel; 60x –0.14 µM/pixel. Dendrites from the *Stratum oriens, Stratum radiatum, and Stratum lacunosum moleculare* regions were analyzed separately. Multiple secondary/tertiary dendrites were analyzed per neuron and the resulting values were averaged. Image analysis was conducted using NIS Elements, and Adobe Photoshop for figure purposes. Brightness was adjusted uniformly across all pixels for a given experiment for figure visualization purposes.

### Statistical analyses

The experimenters were blinded to the examined conditions while carrying out experiments and analyzing data. Bar graphs are depicted as means ± SEM, the distribution of biological replicates (means of culture / individual mice) and – in the case of culture experiments – values for individual synapses / neurons are depicted on the right of each bar as large or small dots, respectively. Statistics were mostly done using a paired t-test on culture means, for electrophysiological recordings and in vivo experiments a Mann-Whitney test on values of individual neurons or mice was performed. *: p < 0.05. **: p < 0.01. ***: p < 0.001.

## ACKNOWLEDGEMENTS

We thank Connie H Wong for thoroughly proofreading the manuscript. This study was supported by grants from the NIMH (R01MH126929 to T.C.S., R00-MH117235 to R.C.S.), the Sloan Research Fellowship (Alfred P. Sloan Foundation) to R.C.S. and the Walter Benjamin Fellowship 505070089 (Deutsche Forschungsgemeinschaft) to D.M..

## AUTHOR CONTRIBUTIONS

D.M., J.M.L., R.C.S., and T.C.S. designed, and D.M., J.M.L. and R.C.S. conducted all experiments. D.M. and R.C.S. validated the conditional knock-in/knock-out mouse lines, D.M. performed all immunocytochemistry, STED/confocal microscopy, and electrophysiological recordings on primary neurons, J.M.L and R.C.S. performed all patching, immunohistochemistry and imaging on brain slices. D.M., J.M.L., R.C.S., and T.C.S. analyzed the data and wrote the manuscript; all authors reviewed and approved the final manuscript.

## CONFLICT OF INTEREST

The authors declare no conflict of interest.

## SUPPLEMENTARY FIGURES AND FIGURE LEGENDS

**Supplementary Figure 1:**
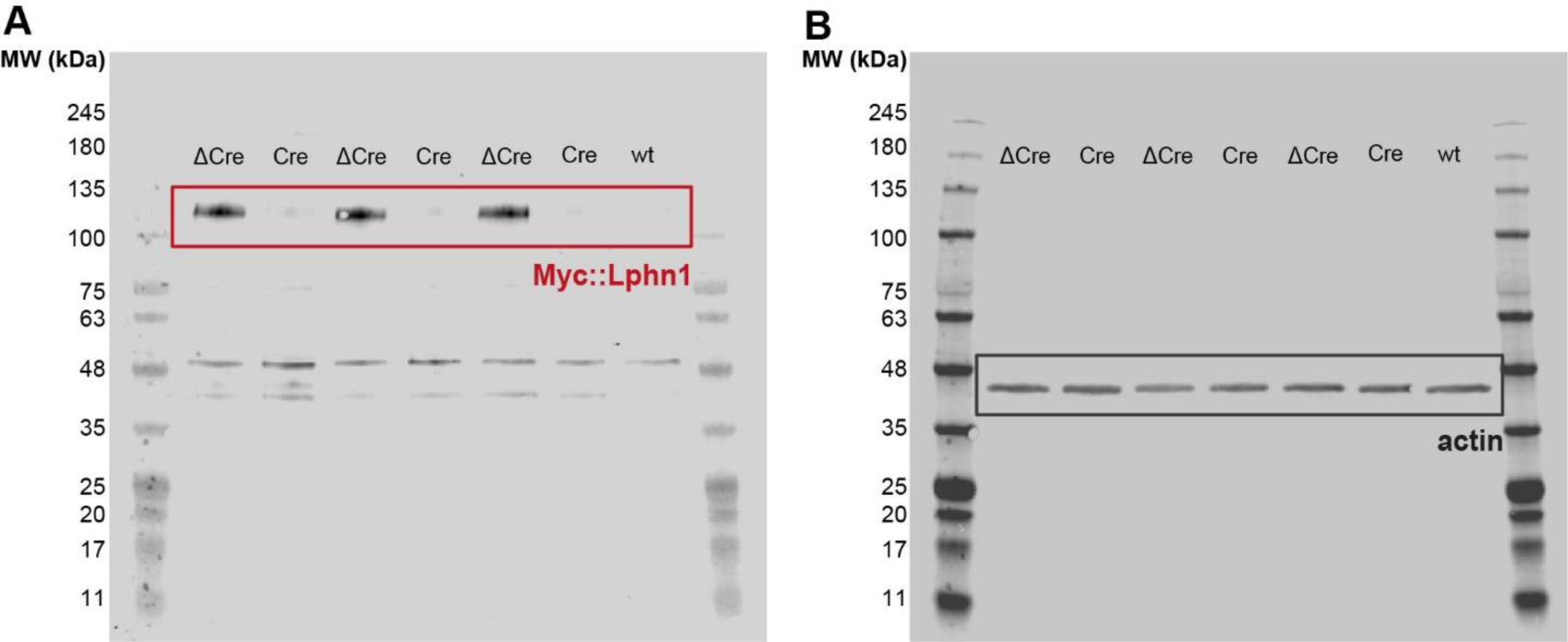
Full immunoblot images for the Lphn1 myc-tag epitope knockin/conditional knockout validation. (A, B) Full images of immunoblots shown in Fig. 1B. myc-tag (A) and actin staining (B) were done with distinct secondary antibodies on the same membrane. Staining for the myc tag shows a specific band at ∼115 kDa in the ΔCre condition but not in the Cre condition and the wild-type (wt) control, corresponding to the cleaved Lphn1 N terminus. Staining for actin (∼40 kDa) shows equal amount of protein has been loaded in all conditions. Unspecific bands are observed with the myc-tag immunoblotting that are not changed by the expression of Cre recombinase and are also present in wild-type controls.

**Supplementary Figure 2:**
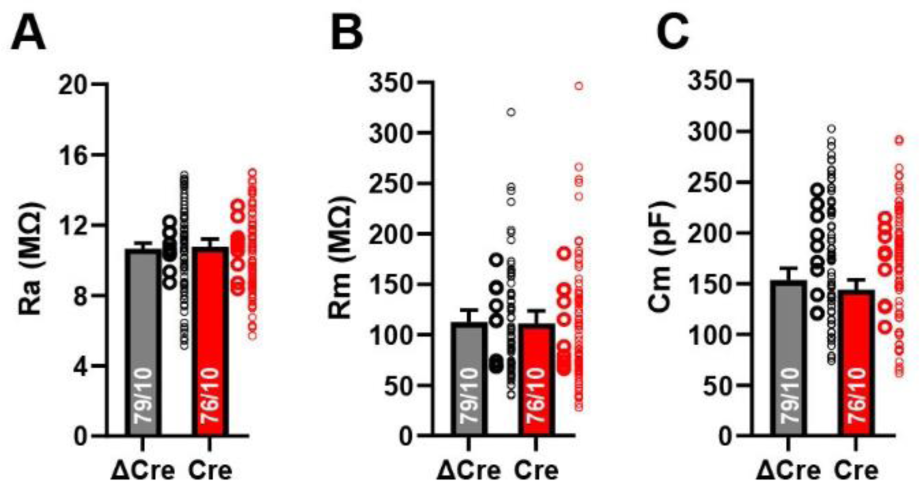
Intrinsic membrane properties monitored during electrophysiological recordings are unchanged by the Lphn1 deletion. Summary graphs show the mean ± SEM access resistance (A), membrane input resistance (B) and membrane capacitance (C). Numbers in bars show the number of neurons/cultures analyzed. Large and small circles next to bars indicate the mean values of each culture and the individual cell values, respectively. Statistics were done using a Mann-Whitney test on individual neuron recordings, but no significant differences have been found.

**Supplementary Figure 3:**
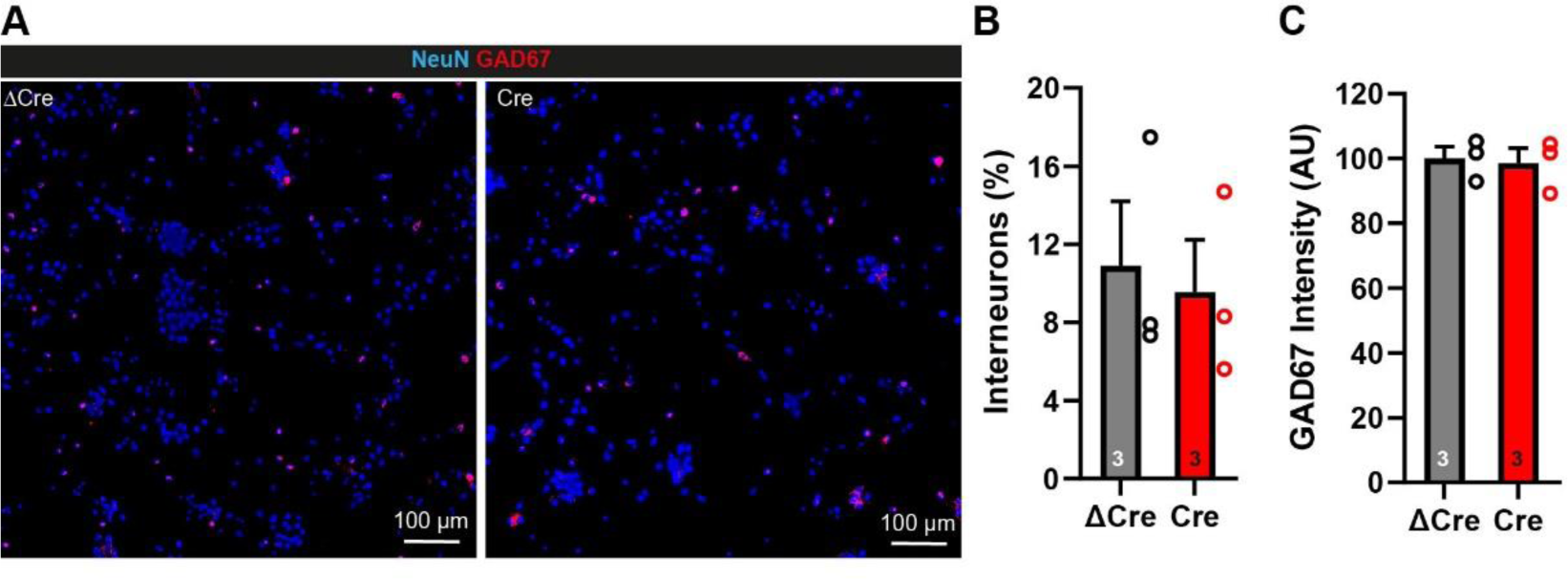
Inhibitory interneuron density is unchanged by the Lphn1 deletion. (A) Representative images of hippocampal neurons cultured from Lphn1 cKO mice. Neurons were infected with lentiviruses expressing active Cre recombinase or inactive mutant ΔCre and immunostained for GAD67 as a marker of inhibitory interneurons and for NeuN as a marker of all neurons. The imaging threshold was set to exclude synaptic GAD67 fluorescence, leaving only somatic fluorescence of interneurons to be analyzed. (B) Quantification of GAD67+ cells (interneurons) as a percentage of NeuN+ cells reveals similar interneuron densities in Lphn1-deficient cultures compared to the control condition. (C) GAD67 staining intensity is unchanged between Cre and ΔCre expressing neurons. Multiple coverslips were scanned and pooled for each condition in each culture (n = 3). Summary graphs show means ± SEMs; circles indicate the mean values obtained in a given independent culture. Statistics were done using paired t-tests, but no significant difference has been found.

## Notes

### Competing Interest Statement

The authors have declared no competing interest.

